# Structure of the full kinetoplastids mitoribosome and insight on its large subunit maturation

**DOI:** 10.1101/2020.05.02.073890

**Authors:** Heddy Soufari, Florent Waltz, Camila Parrot, Stéphanie Durrieu, Anthony Bochler, Lauriane Kuhn, Marie Sissler, Yaser Hashem

## Abstract

Kinetoplastids are unicellular eukaryotic parasites responsible for human pathologies such as Chagas disease, sleeping sickness or Leishmaniasis^1^. They possess a single large mitochondrion, essential for the parasite survival^2^. In kinetoplastids mitochondrion, most of the molecular machineries and gene expression processes have significantly diverged and specialized, with an extreme example being their mitochondrial ribosomes^3^. These large complexes are in charge of translating the few essential mRNAs encoded by mitochondrial genomes^4,5^. Structural studies performed in *Trypanosoma brucei* already highlighted the numerous peculiarities of these mitoribosomes and the maturation of their small subunit^3,6^. However, several important aspects mainly related to the large subunit remain elusive, such as the structure and maturation of its ribosomal RNA^3^. Here, we present a cryo-electron microscopy study of the protozoans *Leishmania tarentolae* and *Trypanosoma cruzi* mitoribosomes. For both species, we obtained the structure of their mature mitoribosomes, complete rRNA of the large subunit as well as previously unidentified ribosomal proteins. Most importantly, we introduce the structure of an LSU assembly intermediate in presence of 16 identified maturation factors. These maturation factors act both on the intersubunit and solvent sides of the LSU, where they refold and chemically modify the rRNA and prevent early translation before full maturation of the LSU.

## Introduction

Kinetoplastids are unicellular eukaryotic parasites, causative agents of several human and livestock pathologies^1^. They are potentially lethal, affecting more than 20 million people worldwide^1^. In part due to their parasitic nature, they strongly diverged from other eukaryotic model species. Kinetoplastids evolved to live in and infect a large variety of eukaryotic organisms in very different molecular environments. Consequently, beyond the general similarities, kinetoplastids species have diverged evolutionarily from each other and their protein sequence identity can be relatively low^7^. They possess a single large mitochondrion, a crucial component of their cellular architecture, where gene expression machineries have also largely diverged, notably their mitochondrial ribosomes (mitoribosomes)^3–5,8–10^. These sophisticated RNA-proteins complexes translate the few mRNAs still encoded by mitochondrial genomes. The mitoribosomes composition and structure greatly diverged from their bacterial ancestor, with the most extreme case described to date being in fact the kinetoplastids mitoribosomes. With highly reduced rRNAs, and more than 80 supernumerary ribosomal proteins (r-proteins) compared to bacteria, completely reshaping the overall ribosome structure. Recent structural studies performed in *Trypanosoma brucei* have highlighted the particularities of this mitoribosome structure and composition as well as the assembly processes of the small subunit (SSU)^3,6^. However, in spite of the very comprehensive structural characterization of the full *T. brucei* SSU and its maturation, several pivotal aspects related to the LSU remained uncharacterized. For instance, a large portion of the LSU at the intersubunit side, including the whole rRNA peptidyl-transfer centre (PTC) along with several r-proteins where unresolved^3^. Moreover, in contrast to the SSU, nearly nothing is known about the LSU maturation and assembly. More generally, the maturation of the mitoribosomes in all eukaryotic species remains largely underexplored. Here, we present a cryo-EM investigation of the full mature mitoribosomes from two different kinetoplastids, *Leishmania tarentolae* (*L. tarentolae*) and *Trypanosoma cruzi* (*T. cruzi*). Most importantly, we reveal the structure of an assembly intermediate of the LSU displaying unprecedented details on rRNA maturation in these very singular mitoribosomes, some of which can probably be generalized to the maturation of all rRNA.

In order to obtain high-resolution reconstructions of the full kinetoplastids mitoribosomes, we purified mitochondrial vesicles from both *L. tarentolae* and *T. cruzi*, and directly purified the mitoribosomes from sucrose gradient (see Methods). All of our collected fractions were analyzed by nano-LC MS/MS (Extended Data Table 1 and 2) in order to determine their proteomic composition. We collected micrographs from multiple vitrified samples corresponding to different sucrose-gradient density peaks for both species and following image processing we obtained cryo-EM reconstructions of the full mitoribosomes, but also the dissociated SSU. Most importantly, we also derived reconstructions of what appeared to be an assembly intermediate of *L. tarentolae* LSU. After extensive rounds of 2D and 3D classification and refinement we obtained the structure of *L. tarentolae* and *T. cruzi* complete and mature mitoribosomes at 3.9 Å and 6 Å, respectively (Extended Data Figs. 1, 2 and 3). Other notable features include the intersubunit contacts and two distinct rotational states in *T. cruzi*. Similarly to *T. brucei*^3^, our cryo-EM analysis revealed a reconstruction of an early initiation complex from *T. cruzi* at 3.1 and 3.2 Å for the body and the head of the SSU, respectively (Extended Data Fig. 4). Further focused refinement on the LSU, SSU head and SSU body generated reconstructions at 3.6, 3.8 and 4 Å, respectively for *L. tarentolae* (Extended Data Figs. 1 and 2), and 3.7 and 4.5 Å for *T. cruzi* LSU and SSU, respectively (Extended Data Fig. 3). Combined, these reconstructions, along with the mass-spectrometry data allowed to build nearly complete atomic models, with only few protein densities still remaining unidentified.

## General description of mature kinetoplastids mitoribosomes

Even though *T. cruzi* and *L. tarentolae* proteins are of relative modest sequence identity (~70%) for such closely related species^7^, the tertiary structures of the r-proteins and rRNAs as well as the overall structure of the mitoribosomes were nearly entirely identical (Extended Data Fig. 3). Both ribosomes are large complexes mainly formed by proteins (Fig. 1 and Extended Data Fig. 3), 68 in the LSU and 54 in the SSU (Extended Data Table 3), rRNA constituting only 15% of the total mass. The SSU was reconstructed to high resolution and appears similar to what was previously observed^3,6^. We obtained reconstructions of the latter in the context of the full ribosomes from *L. tarentolae* and *T. cruzi*, but also dissociated in the context of what appears to be a partial initiation complex from *T. cruzi*, bound to mt-IF3, which accumulated naturally in our samples (Fig. 1 and Extended Data Fig. 4). In the presence of mt-IF3, the 9S rRNA, in particular the highly reduced helix 44, and uS12m are clearly stabilized. However, in the context of the full ribosome, mt-IF3 is no longer present and the 9S rRNA along with uS12m at the subunit interface are significantly more flexible resulting in a scanter resolution. Moreover, for the *T. cruzi* mitoribosome reconstruction, two distinct rotational states are observed, strongly suggesting a fully assembled mitoribosome structure capable of undergoing different conformational states (Extended Data Fig. 3). In contrast to what is observed in prokaryotes and other mitoribosomes, very few of the intersubunit bridges are strictly conserved and most of them rely on protein-protein interactions. However, most of the observed intersubunit bridges are spatially conserved as compared to other known ribosomes but involve here kinetoplastid-specific r-proteins. (Extended Data Fig. 5).

**Figure 1.**
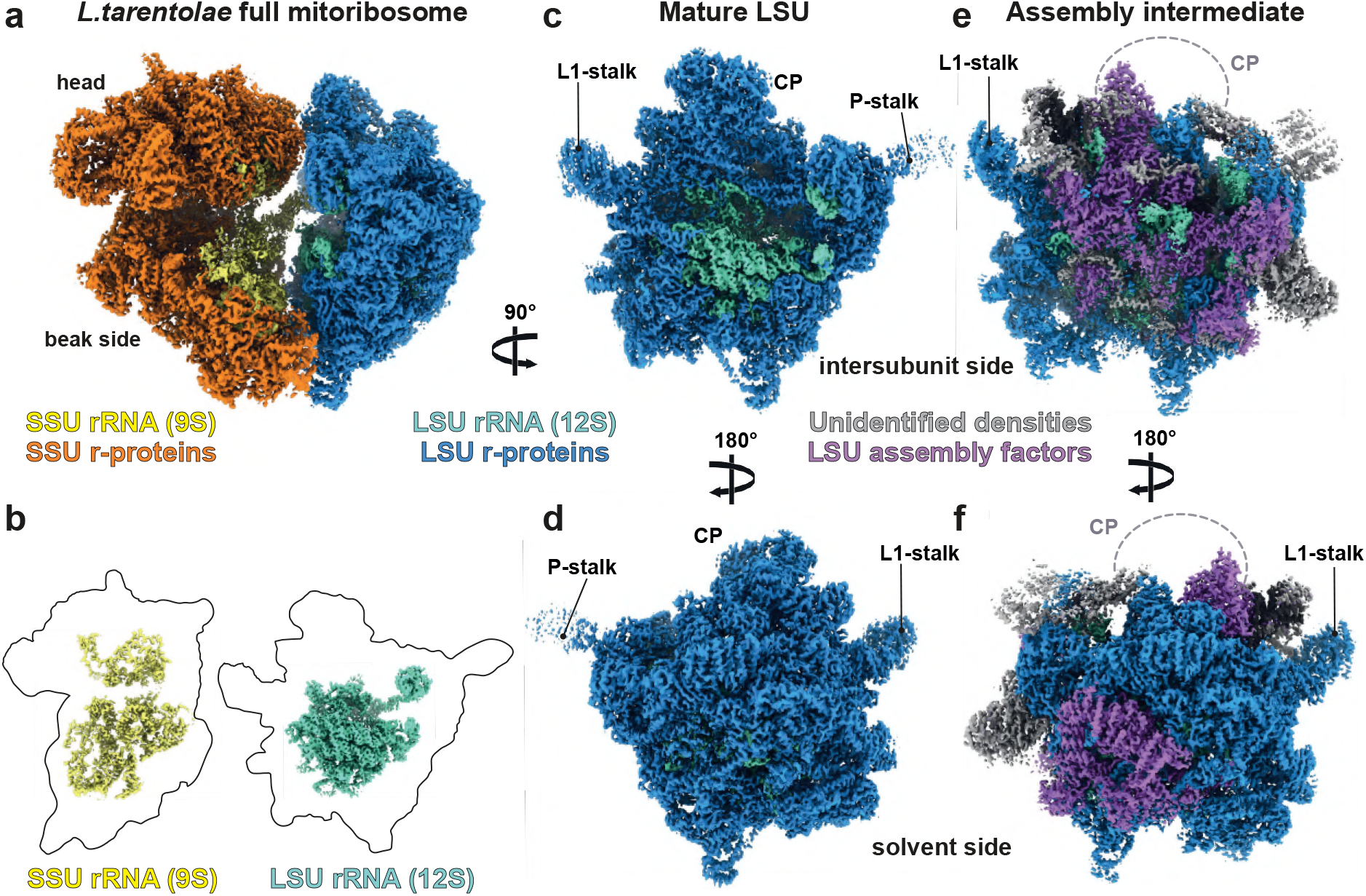
Cryo-EM reconstructions of the mature and assembly intermediate *L. tarentolae* mitoribosome. (**a**) Composite cryo-EM map of the full and mature mitoribosome of *L. tarentolae*. SSU is depicted in orange shades, with proteins in orange and rRNA in yellow and LSU is depicted in blue, with proteins in dark blue and rRNA in aquamarine. (**b**) Relative sizes of the rRNA (colored segmented densities) compared to that of the r-proteins (presented as thick black contours). (**c** and **d**) present the cryo-EM reconstructions of the mature LSU viewed from the intersubunit (**c**) and solvent (**d**) side, compared with the LSU assembly intermediate subunit (**e-f**). Densities corresponding to assembly factors are shown in purple, grey correspond to unidentified factors.

The reconstruction of the large subunit derived from the full mature ribosome (Fig. 1) allowed us to build the entire 12S rRNA including domains IV and V at the interface subunit (Figs. 1 b-c and 4), revealing the catalytic PTC, as well as its intertwined interactions with r-proteins. The structure of the domains IV and V is globally conserved, even when compared to classical prokaryotic ribosomes (Fig. 4). Among the 68 LSU characterized proteins, uL2m, uL14m, and extensions of bL19m, mL68, previously unaccounted for^3^, were visualised and detected by mass-spectrometry (Fig. 2). These proteins and extensions squeeze through the rRNA and stabilize it in a very intricate manner, compared to the classical prokaryotic and cytosolic ribosomes. Surprisingly, compared to what was previously described on the solvent side of the LSU in *T. brucei*^3^, proteins mL67, mL71, mL77, mL78 and mL81, localized close to the peptide channel exit, are absent (Fig. 1 and 2). However, these proteins are present in the assembly intermediate, as described and discussed hereafter.

**Figure 2.**
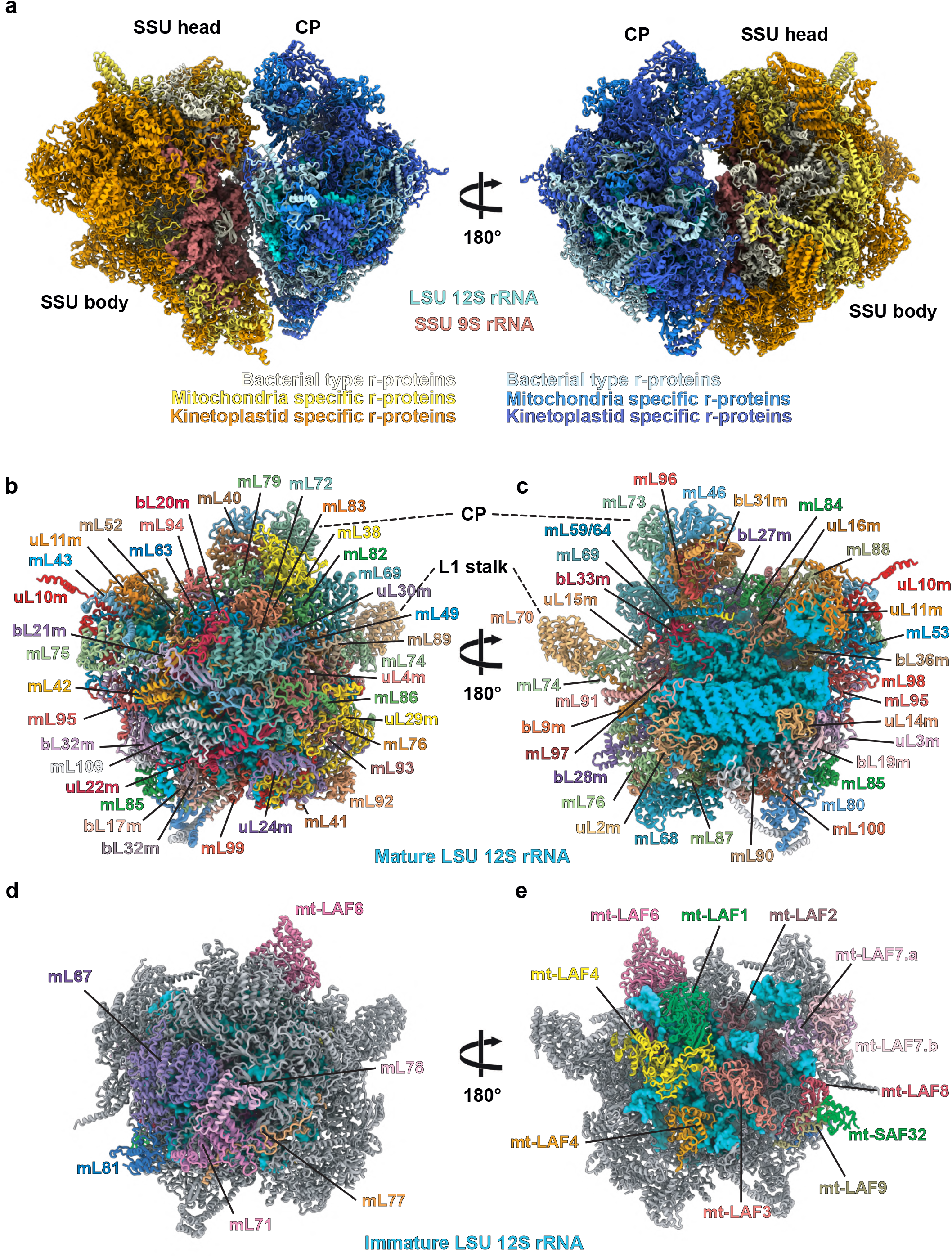
Atomic models of *L. tarentolae* mitoribosome and LSU assembly intermediate. (**a**) Overall model of *L. tarentolae* mitoribosome. 12S rRNA is colored in cyan (LSU) and the 9S rRNA in brick (SSU). Ribosomal proteins are colored in shades of blue (LSU) and shades of yellow (SSU), according to their conservation. Solvent side (**b**) and intersubunit side (**c**) of the mature LSU with each individual r-proteins annotated and displayed with a different color. Solvent side (**d**) and intersubunit side (**e**) of the LSU assembly intermediate. Conserved proteins in the mature LSU (**b-c**) are depicted in gray and assembly factors are colored in different shades and individually annotated.

## General description of the LSU assembly intermediate

During *L. tarentolae* 3D classification, a class of particles presenting LSU-like features was found to naturally accumulate in our sample (Extended Data Fig. 1). This class is rather distinct from classical LSU classes observed previously. After refinement and post-processing, this class resulted in a 3.4 Å reconstruction of the complex (Fig. 1). This reconstruction, when compared to the mature LSU, allowed us to visualize significant differences. Indeed, a large portion of the 12S rRNA appears to be unfolded, or at least highly flexible at the intersubunit face (Fig. 4). In addition, large well resolved protein densities were observed occupying the position of the rRNA at this region. Moreover, the whole central protuberance (CP) is missing and several additional protein densities were observed on the solvent side (Fig. 1 e-f). Therefore, it clearly appeared that this complex corresponds to an LSU assembly intermediate. Our cryo-EM reconstructions, along with the MS/MS analysis, allowed us to build an atomic model of this LSU assembly intermediate (Fig. 2 d-e). Compared to the mature LSU, this assembly intermediate lacks 18 core r-proteins but includes 16 maturation factors (including mL67, mL71, mL77, mL78 and mL81 that were previously described as r-proteins^3^). On the interface, domain IV of the 12S rRNA appears unfolded and several portions of the rRNA undergo refolding or modifications.

## Solvent side

On the solvent side, close to the peptide channel exit, a crown of proteins composed of mL67, mL71, mL77, mL78 and mL81 was observed in the assembly intermediate (Fig. 3), but not on the full mature mitoribosome. The analysis of their structure and interaction network can suggest two main roles. Parts of them hold the 3’ and 5’ extremities of the 12S rRNA. Indeed, the 5’ end of the 12S rRNA is surrounded by several positively charged residues of mL67 coordinating and stabilizing this extremity (Fig. 3 a). On the 3’ end, mL67 again, with the help of mL81 as well as r-proteins bL17m and mL85, stabilize this extremity that is more flexible in the mature LSU (Fig. 3 b), as suggested by its scanter density in the latter. Hence, in the context of the assembly intermediate, these proteins, mainly mL67, hold both 12S rRNA extremities, most likely stabilizing the rRNA in its r-protein shell during its maturation.

**Figure 3.**
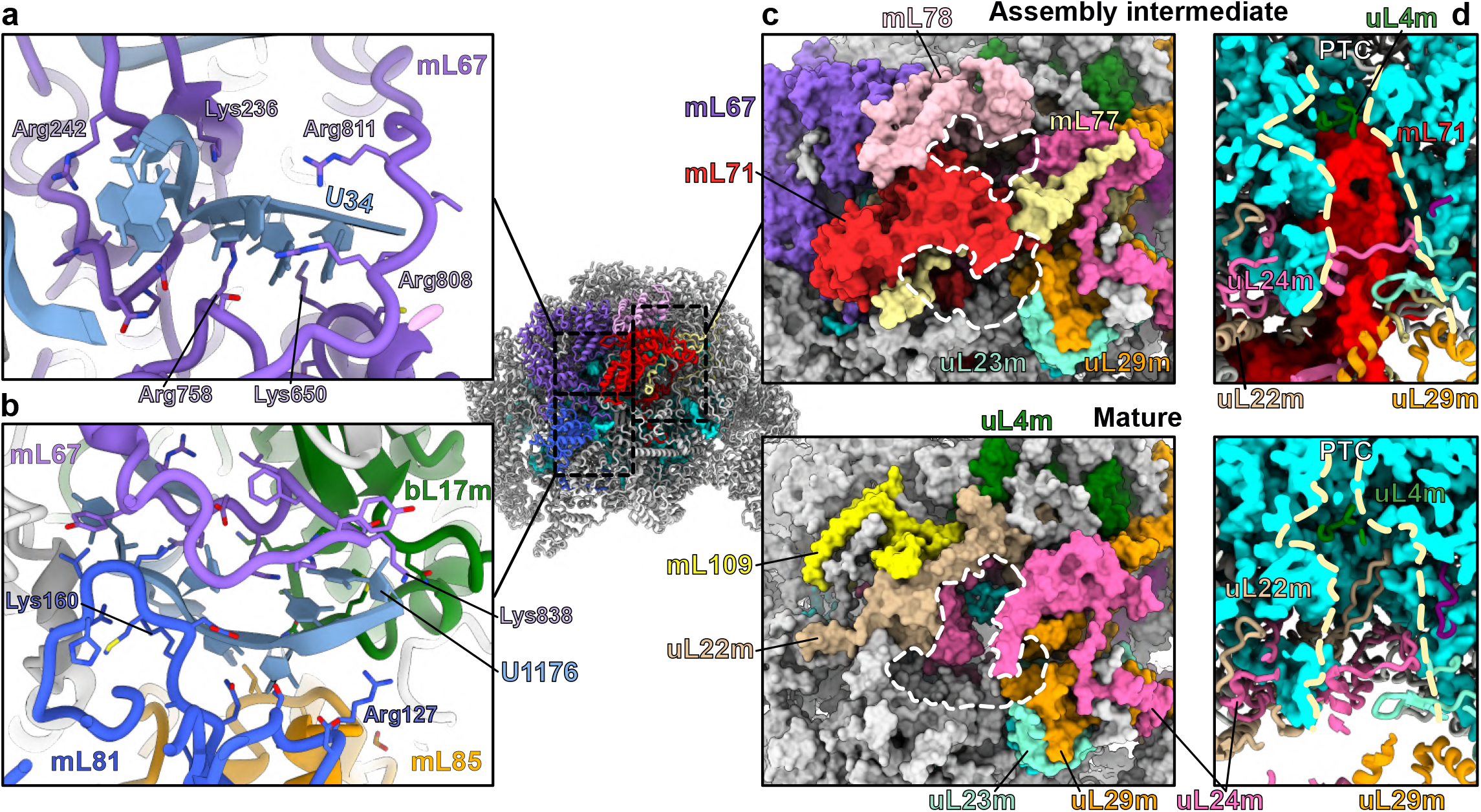
Specific factors hold the 12S rRNA termini and probe the peptide channel exit on the solvent side of the immature LSU. On the solvent side, 5 maturation factors act together to hold the extremities of the rRNA (**a-b**) and probe the peptide exit channel (**c-d**). The 12S 5’ termini (**a**) is held by several positively charged residues of mL67, and the 3’ extremity (**b**) is stabilized by mL67 and mL81 as well as r-proteins mL85 and bL17m. U34 and U1176 indicate the terminal rRNA residues. (**c** and **d**) present the peptide channel of the assembly intermediate (top) and mature (bottom) LSU from top (**c**) and cut view (**d**). Proteins mL71, mL77, mL78 reshape the peptide exit channel by slightly extending and splitting it in two in the assembly intermediate compared to the mature LSU (**c**). Dashed white contours delimit the peptide channel exit orifices in the mature and immature LSU. Maturation factor mL71 completely blocks the exit channel through its N-terminal domain (**d**). Dashed yellow lines delimit the peptide channel in the longitudinal axis.

**Figure 4.**
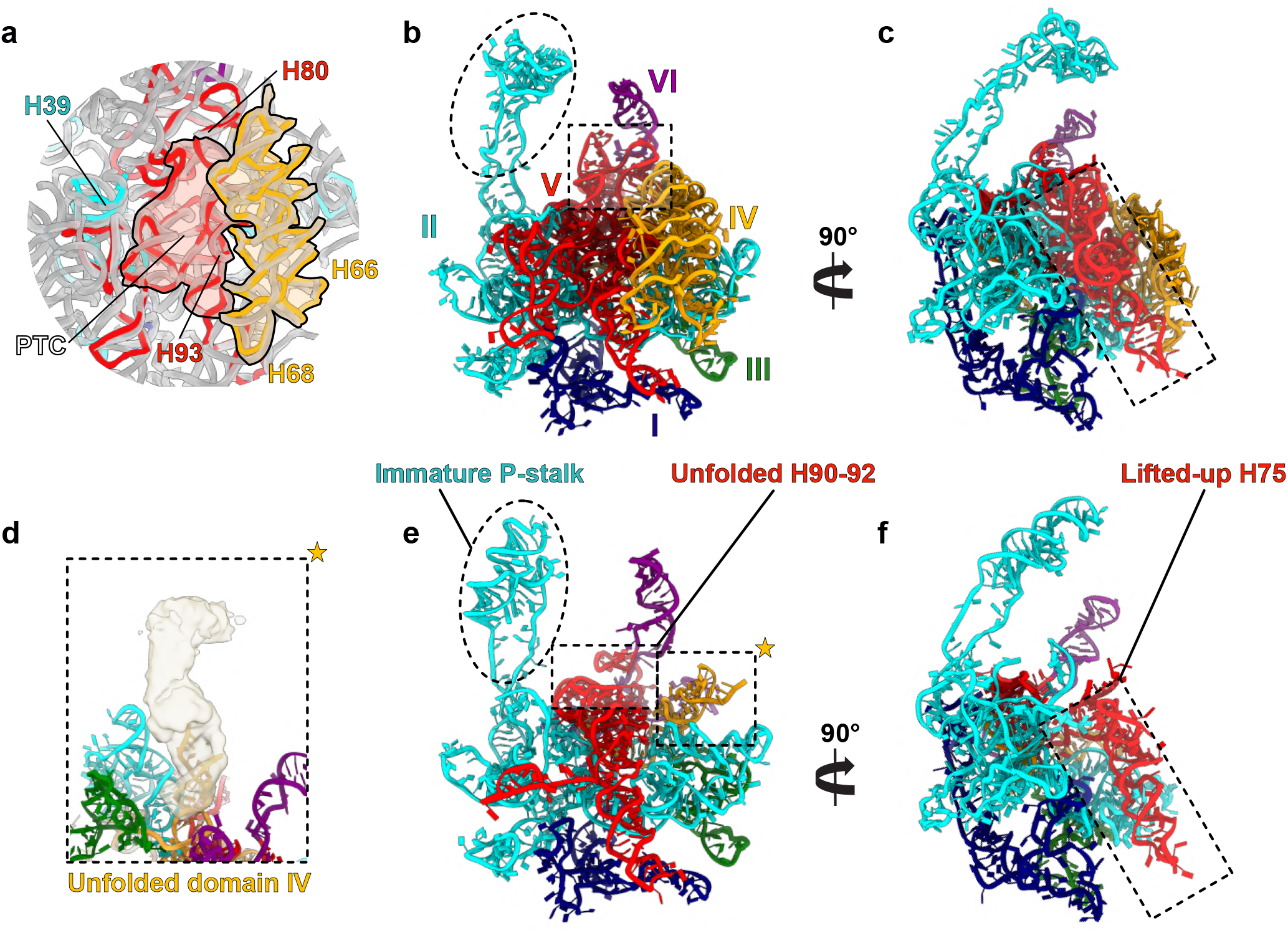
Comparison of the mature and assembly intermediate 12 rRNA. Comparison between mature (**a-c**) and assembly intermediate (**d-f**) 12S rRNA. Each rRNA domains are colored differently. The rRNA is presented from the intersubunit view (**b-e**) and a side view (**c-f**). (**a**) Superimposition of the 12S mature rRNA with *E. coli* 23S rRNA, displayed in gray, shows that the structure of domain IV (orange) and the PTC is highly conserved. Between the mature and assembly intermediate rRNA, the main differences are observed in domain II, where the P-stalk is folded differently, in domain IV which is entirely unfolded (**d**), and domain V which is unfolded from H90 to H92 (**e**) and where H75 is entirely lifted-up (**f**). Most of the maturation factors act on domain V which is accessible thanks to the unfolded domain IV.

Moreover, next to the rRNA extremities, the peptide channel exit is entirely remodeled compared to the mature LSU, as proteins mL71, mL77, mL78 reshape the peptide exit. These proteins slightly extend and split the channel exit in two parts (Fig. 3 c). However, the channel itself is completely blocked by the N-terminal part of mL71, which therefore would prevent active translation (Fig. 3 d), similarly to mL45 that prevents translation until the mitoribosome is bound to the mitochondrial inner membrane in mammals^11^. Upon maturation, these factors are disengaged, and a classical peptide channel is restored, similar in structure to bacterial ribosomes (Fig. 3 c-d). Hence, we hypothesize that these proteins, previously observed in *T. brucei^3^*, are actually maturation factors and that the LSU reconstruction of *T. brucei* is more likely to be a late assembly intermediate (Fig. 7 b). Upon their dissociation from the mature LSU, a novel protein that we termed mL109 replaces mL67 and marks the maturation of this region (Fig. 3).

## Intersubunit side

On the intersubunit side, a large portion of the 12S rRNA, corresponding to domain IV, appears unfolded and several proteins occupy its position, acting on the immature domain V. Based on our cryo-EM reconstructions coupled with MS/MS analysis and structural and sequence homologies, 11 factors were identified bound to the intersubunit side (Fig. 5). We termed these proteins “mt-LAFs” for mitochondrial LSU assembly factors, consistently with the proposed nomenclature^6^. Some of these factors belong to the family of GTPases and RNA helicases/isomerases, classic actors of ribosome biogenesis and were observed with their respective ligands and rRNA targets (Fig. 6 e). Others are RNA modification enzymes. Part of these factors exhibit homology with bacterial and eukaryotic maturation factors previously identified. However, most of these ribosome maturation factors have never been structurally captured in an assembly intermediate context.

**Figure 5.**
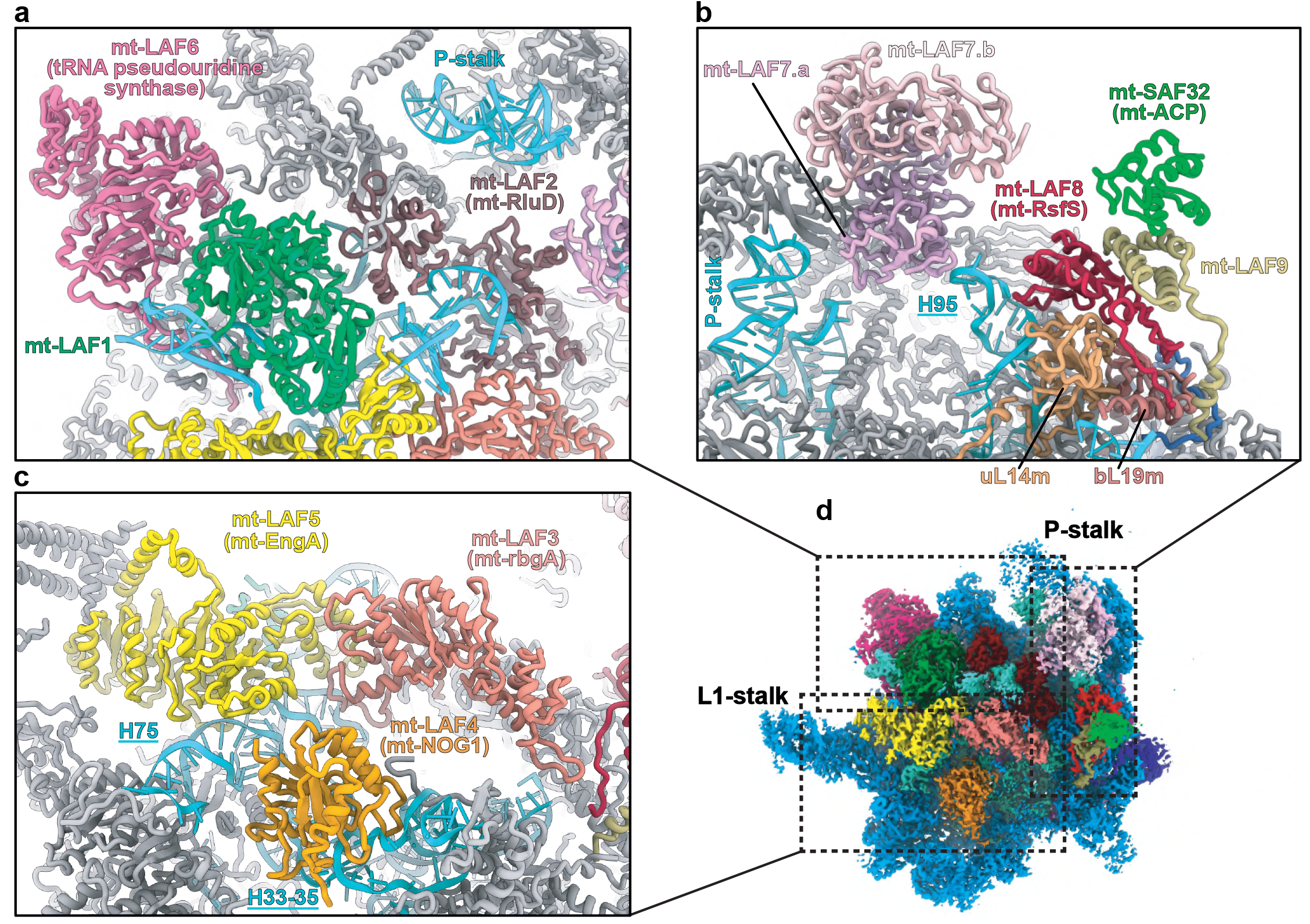
Overall view of the maturation factors on the intersubunit side. On the intersubunit side of the immature LSU, 11 maturation factors act together to remodel and modify the rRNA (**d**). (**a**) mt-LAF6 and mt-LAF1 occupy the position of the central protuberance (CP). mt-LAF1 and mt-LAF2 interact with remodeled rRNA. (**b**) Mt-LAF8 together with mt-LAF9 and mt-SAF32 interact with uL14m and H95 to block the SSU association. (**c**) Arrangement of the GTPase triptych. The GTPases mt-LAF5 and mt-LAF4 bind respectively the rRNA H75 and H37 while mt-LAF3 occupies a central place in the complex and makes a bridge between the GTPases and mt-LAF2 and mt-LAF8.

**Figure 6.**
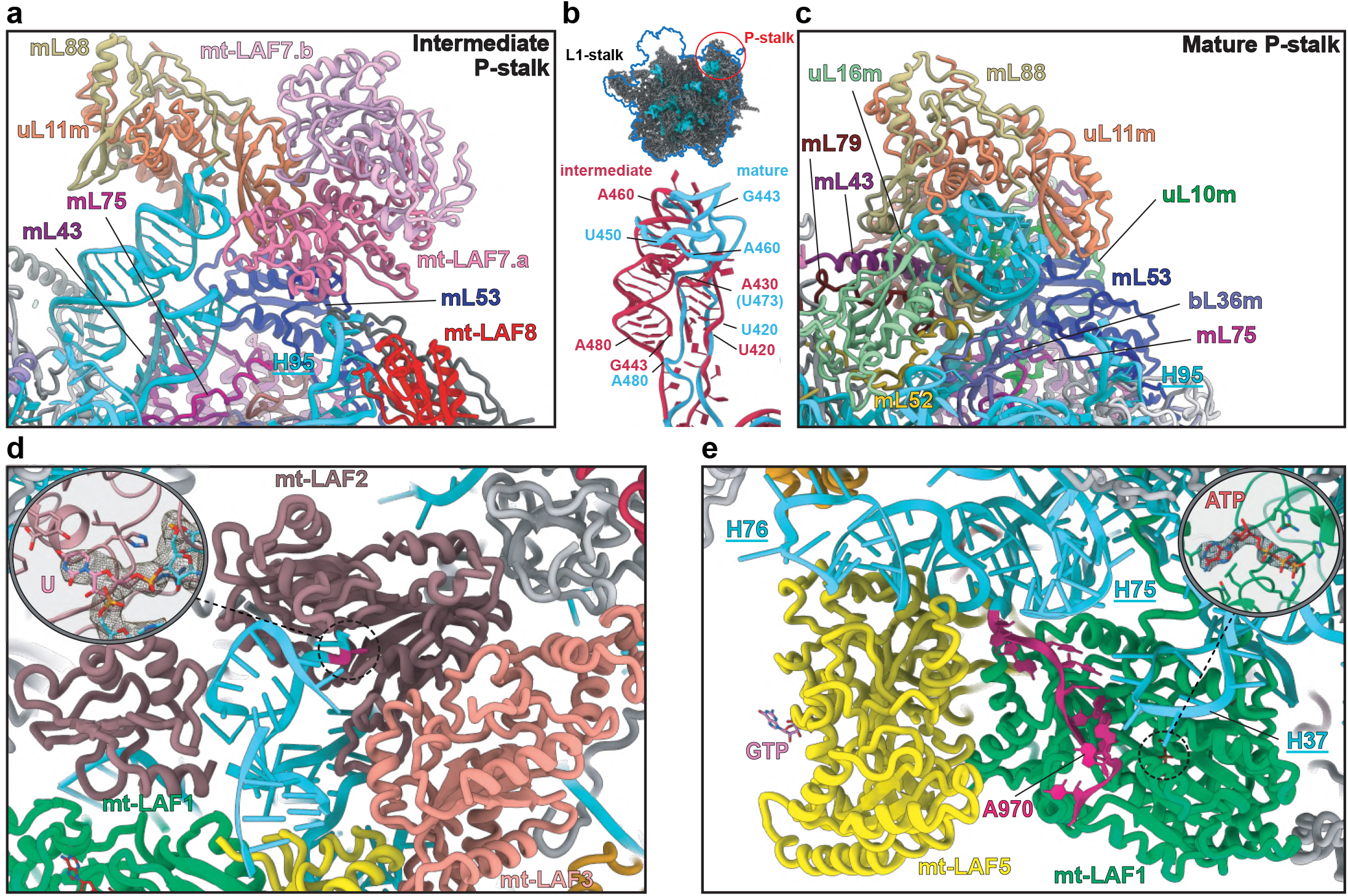
Maturation of the Leishmania mitochondrial LSU rRNA. Details on how maturation factors interact with, and affect the 12S RNA (**a**) P-stalk region of the intermediate LSU. mt-LAF7 homodimer contacts uL11m and mL53, the two helices of the immature P-stalk as well as the helix H95 are adjacent to the dimer. (**b**) Overlap of the two forms of rRNA P-stalk structure, immature in red and mature in blue. (**c**) P-stalk region of the mature LSU. (**d**) Insight into the rRNA modification by the pseudouridine synthase mt-LAF2. In order to be catalyzed the rRNA is refolded into a helix structure, probably to be recognized by mt-LAF2, the uridine subtract is flipped out of the helix and engaged in the catalytic core of the enzyme. (**e**) The two factors mt-LAF5 and mt-LAF1 bind jointly to the rRNA. The dead box helicase mt-LAF1 is in an active state with a single strand RNA of the immature H88, shown in burgundy, in its catalytic core and loaded with an ATP molecule. Doted circle corresponds to magnified region.

The first noticeable difference with the mature LSU is the absence of central protuberance (CP). Indeed, 8 proteins composing the CP are missing and some of the surrounding proteins adopt different conformations (Extended Data Fig. 6). In the assembly intermediate, the binding of the CP block is prevented by mt-LAF1 and mt-LAF6 (Fig. 5 a and Extended Data Fig. 6). The factor mt-LAF1 is a DEAD-box RNA helicase, where both the catalytic and ATP-binding sites are conserved^12^, and the ATP ligand is clearly observed (Fig. 6 e). It occupies the spatial positions of bL27m, bL31m, bL33m and mL82. The role of this family of helicase proteins is to unwind RNA structures and/or RNA-protein complexes, thus they are highly involved in ribosome maturation. Mt-LAF1 shows similarity with Has1, previously described in yeast pre-60S maturation and with Mss116, involved in yeast mitoribosome maturation^13,14^ (Extended Data Fig. 7). Here, it contacts two different segments of the 12S rRNA, the tip of H37 loop (residues 341-346) shifting the whole helix, compared to the mature LSU (Fig. 6e), and a single strand (residues 968-975) that will become part of H88 from domain V after its maturation. Thus, mt-LAF1 appears to be involved in the maturation of domain V rRNA. The second factor, mt-LAF6, is in direct contact with mt-LAF1, but does not contact rRNA. Compared to the mature LSU, it occupies the place of mL38, mL69 and mL82. It shows high homology with tRNA pseudouridine synthases, however several residues of the active site are not conserved (Extended Data Fig. 8), hinting that the protein has lost its original function here. Its role seems to be a placeholder for CP r-proteins, thus acting together with mt-LAF1 to prevent the CP r-proteins premature association. When mt-LAF 1 and 6 are disengaged, the CP r-proteins (composed of 8 proteins) can dock onto the LSU (Extended Data Fig. 6). Moreover, the mL59/64 C-terminal part that is flexible in the assembly intermediate can contact and stabilize the CP in the mature LSU context.

A large portion of the 12S rRNA is unfolded, which corresponds to the entire domain IV (Fig. 4), and several maturation factors are present in the region that will later be occupied by the rRNA after its maturation, including a triptych of GTPases, mt-LAF3, mt-LAF4 and mt-LAF5 (Fig. 5 c and Extended Fig 9). While some of these GTPase have already been identified in bacteria^15,16^ (Extended Data Table 3), this is the first time that they are observed acting together on a maturating ribosomal complex at the reported resolution. GTPases have widespread roles in the cell^17^. These enzymes usually exist in two states, active and inactive, where the transition between the states is triggered by GTP hydrolysis. This results in conformational changes of the enzyme that allow its binding to or release from specific targets. mt-LAF5 has two characteristic G-domains, similarly to its bacterial homologue, EngA, also known as GTPase Der^15^. Its role is crucial for domain IV PTC folding in bacteria, a role that is likely conserved here for mt-LAF5 given that their binding sites are similar. mt-LAF4 is bound to H33-35 and appears here to prevent uL2m association. Mt-LAF3 (homolog to the bacterial GTPase RgbA^18^) is in contact with mt-LAF 5 and 2. However, it is the only GTPase that does not make significant contacts with the rRNA (Fig. 5 c), suggesting that it has either already performed its function, or is still awaiting for its rRNA target (Extended Data Fig. 9 b).

Three RNA modification enzymes were found in the assembly intermediate structure, one pseudouridine synthase (termed “mt-LAF2”) and two methyltransferases forming a homodimer (termed “mt-LAF7 a and b”). The protein mt-LAF2 is a pseudouridine synthase D family enzyme that shows high homology with RluD, a bacterial enzyme responsible for the modification of several residues on H69 in domain IV of 23S rRNA^19,20^ (Extended Data Fig. 8).In our reconstruction, we clearly observe a uridine residue flipped-out of an RNA regular helix to engage the catalytic core of the enzyme (Fig. 6 d). Mt-LAF7 a and b are RNA 2’O-methyltransferases that belong to the SpoU family, which are enzymes wildly involved in methylation of the 23S rRNA in prokaryotes^21,22^ (Extended Data Fig. 10). They are located at the P-stalk region (Fig. 6 a). Surprisingly, the P-stalk structure in the immature rRNA is radically different compared to its mature form. Indeed, the P-stalk rRNA undergoes drastic structural rearrangements between the immature state, where it is shaped into two adjacent large regular helices (Fig. 6 a), and the mature P-stalk (Fig. 6 c), where it doesn’t form any RNA helices. Interestingly, even though the fold is radically different, the immature P-stalk rRNA still spans all the way and interacts with the distal uL11m (Fig. 6 b) that retains its position in the mature P-stalk. While the nearby homodimeric factors mt-LAF7.a/mt-LAF7.b are in close proximity to the rRNA, they cannot be unambiguously interpreted to bind it directly. mt-LAF7.a/mt-LAF7.b contact uL11m and mL53 inducing the shift of the P-stalk r-protein block compared to the mature structure. Moreover, several surrounding r-proteins such as uL10m or uL16m are absent. Mt-LAF7 a and b are most likely the homologs of the NHR SpoU described in *S. actuosus*^21^;. which are involved in an adenine methylation in H43 of the P-stalk. Due to their position in our assembly intermediate, it is possible that they modify the P-stalk rRNA. The latter will thus be reshaped into its mature structure afterwards, probably under the action of a helicase analogue to mt-LAF1 (Extended Data Fig. 7 c).

Lastly, a complex formed by three assembly factors is found bound to uL14m, close to the P-stalk. This complex is formed by mt-LAF8, directly contacting uL14m and the tip of H95 (residues 1133-1136), mt-SAF32 and mt-LAF9 (Fig. 5 b and 7 a). This complex is nearly identical to the MALSU1:L0R8F8:mt-ACP complex observed in the human mitoribosome LSU assembly intermediate^23^. Indeed, each factor shows high homology to their mammalian homologs, mt-LAF9 corresponding to L0R8F8, mt-SAF32 to mt-ACP and finally mt-LAF8 to MALSU1. The latter shows high homology to the conserved and well characterized bacterial ribosome silencing factor (RsfS) (Fig. 7 e). RsfS is known to prevent premature subunit association during diminished nutrient availability, a molecular mechanism mimicked by eIF6 in eukaryotes^24,25^. However, in mitoribosomes, as shown in human, the RsfS homolog alone is not able to prevent subunit association and the anti-subunit association activity can only be achieved in the presence of mt-SAF32 and mt-LAF9.

**Figure 7.**
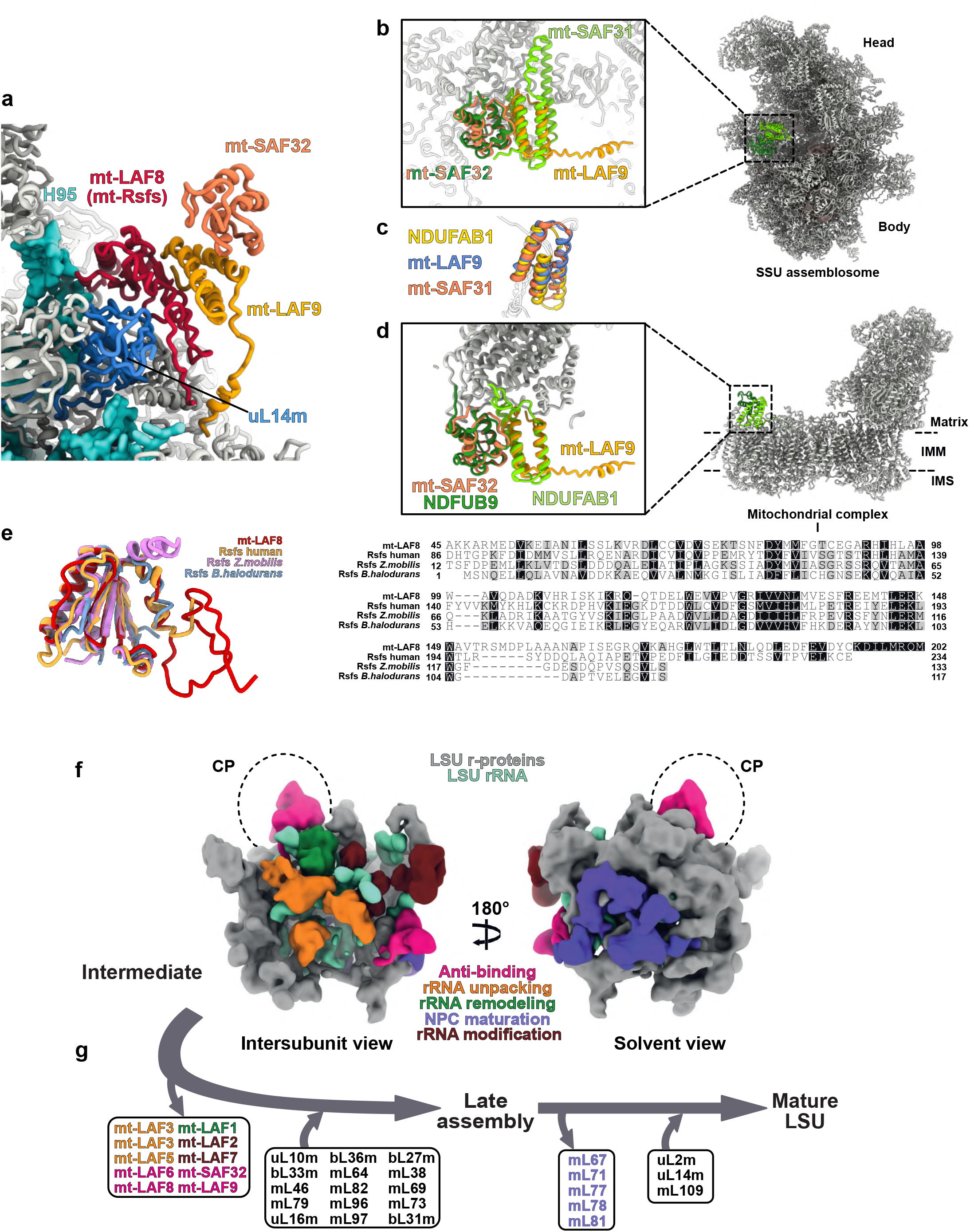
Key factors and schematic view of the stepwise assembly. Zoom on the mt-LAF8:mt-LAF9:mt-SAF32 factors involved in anti-subunit association (**a**). mt-LAF8 is the homolog of the bacterial protein Rsfs (**e**) and mt-SAF32 and mt-LAF9 are also found in mitochondrial respiratory complex I as supernumerary proteins. In (**b**) mt-SAF32 and mt-LAF9 are superimposed with mt-SAF32 and mt-SAF31 found in the SSU assemblosome of *T. brucei*^6^ (PDB: 6SGB). In (**d**) mt-SAF32 and mt-LAF9 are respectively superimposed with NDUFB9 and NDUFAB1, here from the yeast mitochondrial complex I^32^ (PDB: 6RFR), homolog proteins are annotated using the human nomenclature of complex I proteins. mt-LAF9, NDUFAB1, and mt-SAF31 are LYR proteins characterized by the LYR-protein fold (three antiparallel helices) shown in (**c**). IMM stands for inner mitochondrial membrane and IMS for inter membrane space. In (**e**), the high structural conservation of Leishmania mt-LAF8 is illustrated by structural superimposition and sequence alignment with the human mitochondrial Rsfs (PDB: 5OOL) as well as two bacterial Rsfs (PDB: 3UPS and 2O5A). (**f**) Schematic view of the LSU assembly intermediate of *L. tarentolae* presented from the intersubunit and solvent sides. Maturation factors are colored by function. Starting from the LSU assembly intermediate in (**f**), (**g**) details the dissociation of maturation factors and the association of r-proteins, considering the *T brucei* LSU^3^ as a late assembly intermediate. Maturation factors are colored as in (**f**) and r-proteins are shown in black.

## Discussion

We describe the structures of the full kinetoplastids mitoribosomes from *L. tarentolae* and *T. cruzi*. These reconstructions allowed us to characterize the complete 12S rRNA of the large subunit revealing notably the yet uncharacterized domain IV and V including the PTC. While kinetoplastids mitoribosomes represent one of the most extreme cases of rRNA reduction^6,26^, these regions are structurally conserved compared to bacterial and other mitochondrial ribosomes (Fig. 4). These rRNA domains represent part of the conserved functional core of the ribosome, where the peptidyl transfer is catalyzed, in addition of being a conserved binding surface to the small subunit^27^.

On the small ribosomal subunit, the 9S rRNA appears to be more flexible than its counterpart LSU 12S rRNA, as indicated by its scanter densities at the intersubunit face (Fig. 1). It is important to highlight that the entire SSU 9S rRNA only conserves a reduced number of secondary structures (10 helices^3^) and most of its nucleotides are organized in single strands that are intertwined with the r-proteins. In our study, its structure was derived from the SSU/mt-IF3 complex where the 9S rRNA densities are solid and interpretable, corroborated by previous studies^3,6^. The reason of the accumulation of these SSU/mt-IF3 complexes is unclear but could reflect a relative low level of translation initiation in *T. cruzi*, at least at the growth stage at which the parasites were harvested. An additional explanation could be the fairly high hydrophobicity index of uS12m that is one of the only two r-proteins (along with uS3m) encoded by the mitochondrial genome in kinetoplastids. Indeed, recruiting such a hydrophobic r-protein to the negatively charged rRNA surface may require the presence of mt-IF3 that stabilizes it on the SSU intersubunit face rRNA (Extended Data Fig. 4). In such case, these SSU/mt-IF3 complexes would correspond to a small subunit ready to initiate translation. The biological rational for the presence in kinetoplastids mitoribosomes of such a hydrophobic r-protein (uS12m) embedded in the rRNA at this strategic region remains to be unveiled but could be related to the regulation of the nuclear-mitochondrial crosstalk.

We also describe an assembly intermediate of the large subunit of *L. tarentolae* mitoribosome. Assembly intermediates in mitoribosomes have only been structurally characterized in mammals for the LSU^23^ and in *T. brucei* for the SSU^6^. Here, our reconstruction enabled the identification of 16 proteins factors involved in various ways in the LSU maturation. A portion of these factors appears to be conserved in bacteria while the rest are specific to kinetoplastids mitoribosome. The kinetoplastid-specific factors are found on a region of the LSU that has evolutionally diverged the most in terms of structure, i.e. on the solvent-side of the LSU. The positions and interactions of these factors suggest two main roles. First, they maintain the partially unfolded and maturing rRNA 3’ and 5’ extremities in the LSU protein shell till the complete maturation of the 12S rRNA (Fig. 3). Second, they prevent any premature translation by blocking the peptide channel exit. This molecular mechanism is a known feature of immature LSU/ribosome. For instance, during cytosolic 60S maturation the Rei1 protein probes and clots the nascent peptide tunnel^28^. Another example can be found in the human mitoribosome where mL45 similarly blocks the channel till the binding to the inner mitochondrial membrane^11^.

The other maturation factors have homologs found in bacterial, cytosolic and mitochondrial ribosome biogenesis processes. They act on the intersubunit side of the LSU where the 12S rRNA is mostly conserved and to some extent similar to bacteria. Amongst these conserved factors, three modification enzymes, two 2’O-methyltransferase (mt-LAF7 a and b) and one pseudouridine synthase (mt-LAF2). The global structures and active sites of mt-LAF 7 and 2 are conserved (Extended Data Fig. 8 and 10). Mt-LAF7 a and b are RNA 2’O-methyltransferases that belong to the SpoU family. Compared to the bacterial enzymes, their structure as well as their dimerization are conserved. Their catalytic core, involved in SAM binding, is conserved as well as their N-terminal domain, involved in dsRNA binding (Extended Data Fig. 10). In the assembly intermediate, mt-LAF7 a and b are located near to three possible targets, H95 and the two helices of the immature P-stalk. Several of these enzymes are involved in methylation of the 23S rRNA in prokaryotes^21,22^.The NHR SpoU described in *S. actuosus*^21^, are involved in an adenine methylation in H43 of the P-stalk. Given that mt-LAF7 a and b are located close to the P-stalk, it is possible that their target is similar to NHR SpoU. The folding of the immature P-stalk in two helices would therefore be crucial for mt-LAF7 a and b to recognize their target and perform methylation. The protein mt-LAF2 is a pseudouridine synthase D family enzyme that shows high homology to RluD, a conserved bacterial enzyme responsible for the pseudouridylation of several residues on H69 in domain IV of 23S rRNA^19,20^. This enzyme is highly conserved in all Rlu bacterial enzymes (Extended Data Fig. 8) and is essential for ribosome biogenesis and cell growth in bacteria^29^. In our reconstruction, we clearly observe a uridine residue flipped-out of an RNA regular helix to engage the catalytic core of the enzyme as previously characterized for these enzymes (Fig. 6 d and Extended Data Fig. 8), strongly suggesting that this factor is catalytically functional.

To date, only few examples of rRNA modifications were characterized in mitoribosomes, the best being described in human^30^. Only three types of modification were identified in the human mitoribosomal rRNA: nucleobase methylation, 2’-O-methylation and pseudouridylation. Given the conservation level of mt-LAF 7 and 2 (Extended Data Fig. 8 and 10) as well as the rRNA regions that they target, it is expected that other modifications found in bacteria^31^ and mitoribosomes^30^ in domains IV and V might also be conserved in kinetoplastids mitoribosomes (such as Um2552 in *E. coli*, equivalent to mitoribosome human Um1369^30^). In our reconstruction, we also observed the presence of an inactive pseudouridine synthase, mt-LAF6, as deduced by the lack of conservation of critical catalytic amino acids, compared to bacterial, human and yeast homologs (Extended Data Fig. 7 a). Its location on the LSU assembly intermediate suggests its repurposing as an anti-binding factor, as it interferes with the binding of the central protuberance (Extended Data Fig. 6).

Three GTPases, all homologs of bacterial proteins, are also found in our reconstruction (Extended Data Fig. 9). The bacterial homologs of two of them, EngA (homolog of mt-LAF5) and RbgA (homolog of mt-LAF3) were previously described in context of the 50S^15,16^. Mt-LAF5, in the GTP state, (Extended Data Fig. 9 a) interacts with H74-75 of domain V and mt-LAF3 that contacts mt-LAF5 through its N-terminal domain. Similarly to the described roles of these proteins in bacteria, they bind the assembly intermediate rRNA to facilitate the folding of domain IV and V helices^15,16^. The third GTPase, mt-LAF4, is homologous to EngB/YihA in bacteria that was structurally characterized alone, outside the context of the maturing LSU. Here, we show that in the *L. tarentolae* LSU assembly intermediate, mt-LAF4 contacts helices H33-35 of domain II and occupies the place of uL2m (Extended Data Fig. 9 c). Finally, the DEAD-box RNA helicase mt-LAF1 is found on the immature H88 (Extended Data Fig. 7). Comparison with other structures of DEAD-box enzymes revealed that, aside from the presence of a flexible insertion domain between the DEAD and helicase domains, the overall organization of the domains is conserved. Ligand and RNA binding are also performed in a conserved fashion. mt-LAF1 is in the closed state of its cycle, acting on a single stranded portion of H88, confirming its activity. Although the role of the flexible insertion domain between the DEAD and helicase domains is unknown, it does not appear to hamper its function. This factor has no obvious bacterial homologs but shows similarity to Has1^14^ and Mss116^13^. While Has1 participates in the maturation of H16 in the pre-60S cytosolic ribosome, Mss116 appears to be mt-LAF1 yeast mitochondrial homolog. The structure of Mss116 was solved alone, off the context of the maturing yeast mitochondrial ribosome^13^. Therefore, our structure provides invaluable insight on the molecular basis of the maturation function of this protein.

Lastly, the mt-LAF8:mt-LAF9:mt-SAF32 complex is composed by the bacteria-conserved Rsfs factor (mt-LAF8) and two repurposed respiratory complex I components (mt-SAF32 and mt-LAF9)^32–34^. The latter are supernumerary subunits of the complex I in eukaryotes, i.e. that they are not part of the core proteins found in bacterial complex I^35^, but were acquired during respiratory complex I evolution in mitochondria^36,37^. Interestingly, they interact together in the same fashion as in the mitochondrial complex I (Fig. 7 a-e). Moreover, in the *T. brucei* SSU assemblosome^6^, mt-SAF32 binds mt-SAF31 that possesses a LYR-domain (LYR designate the conserved tripeptide motif Leucine, Tyrosine and Arginine). LYR-domains are characterized by a specific fold made of three antiparallel helices. Mt-LAF9 found in our structure also contains a LYR-domain, thus mt-SAF32 most likely recognize the characteristic LYR fold in order to bind to mt-SAF31 and mt-LAF9 in the context of SSU and LSU assembly intermediates. Even though respiratory chain components in kinetoplastids appear strongly divergent from the rest of eukaryotes^38^, the presence of the mt-LAF9:mt-SAF32 in both human LSU assembly intermediate and complex I, in addition to the presence of mt-SAF32 in kinetoplastids SSU assembly intermediate, suggests a multi-role of these proteins and hint at a possible crosstalk between mitoribosome maturation and respiratory complex assembly.The natural accumulation of the LSU assembly intermediate in our sample could be promoted by the presence of the mt-LAF8:mt-SAF32:mt-LAF9 complex, which might be stress-induced or caused by diminished nutrient availability, likewise in bacteria. Interestingly, as its name already indicates, mt-SAF-32 is also found in the SSU assembly intermediates, likewise at a peripheral position of the intersubunit side^6^.

In conclusion, our cryo-EM structures of kinetoplastids mitoribosomes, especially the yet uncharacterized LSU assembly intermediate, shed new lights into ribosome maturation. Even though several proteins are specific to kinetoplastids, most of the maturation factors present in our structure have homologs in other eukaryotes, mitochondria or bacteria (Extended Data Table 3). Hence, our structure unveils the mode of actions of these factors, for which no high-resolution structures in native conditions existed. It also highlights that despite being highly divergent, kinetoplastids mitoribosomes retained several canonical maturation pathways for their rRNAs and probably follow an analogous mechanistic pattern (Fig. 7 f-g). Indeed, in concert, a set of factors intercalate with the premature rRNA (GTPases) to mechanically unpack different premature rRNA domains, allowing RNA helicases to disentangle and unwind the rRNA and several families of modification enzymes to access their targets. The remodeling of the rRNA is also aided by the recruitment of the r-proteins early during the maturation process. This is especially true in the kinetoplastids mitochondria, where most of the r-proteins are already recruited at the observed stage and form a thick protein shell holding the rRNA together. At the same time, various anti-association factors prevent premature binding of r-proteins and the SSU joining.

In spite of large environmental and host diversities in which these parasites evolve, their mitoribosomes are conserved structurally among kinetoplastids. Therefore, our structures could serve as a basis for future experiments to develop more effective and safer kinetoplastid-specific therapeutic strategies. Finally, as kinetoplastids undergo extensive mitochondrial reorganization^39^ during their life cycle, both in vectors and hosts, it would also be interesting to study mitoribosomes from different stages of development from the parasite’s life cycle.

## METHODS

Methods and any associated references are available in the online version of the paper.

## ACKNOWLEDGEMENTS

This work has benefitted from the facilities and expertise of the Biophysical and Structural Chemistry platform (BPCS) at IECB, CNRS UMS3033, Inserm US001, Bordeaux University. We thank A. Bezault for assistance with the Talos Arctica electron microscope. We thank J. Chicher and P. Hamman of the Strasbourg Esplanade proteomic analysis for the proteomic analysis.

This work was funded by a European Research Council Starting Grant (TransTryp ID:759120) and the IdEx junior excellence chair program of the Université de Bordeaux to YH and the French National Program “Investissement d’Avenir” (Labex MitoCross), administered by the “Agence National de la Recherche”, and referenced [ANR-11-LABX-0057_MITOCROSS] to MS. This project was supported by “Institut National de la Santé et de la Recherche Médicale” (INSERM), “Centre National de la Recherche Scientifique” (CNRS) and Université de Bordeaux.

## AUTHOR CONTRIBUTIONS

YH, HS and FW designed and coordinated the experiments. CP and SD purified kinetoplastids mitochondria. FW purified mitoribosomal complexes. LK performed the mass-spectrometry experiments. HS acquired the cryo-EM data and processed the cryo-EM results with YH and FW. HS, YH, FW and AB built the atomic models. FW, HS, MS and YH interpreted the structures. FW, HS, CP, YH and MS wrote and edited the manuscript. YH directed the research.

## COMPETING FINANCIAL INTERESTS

The authors declare no competing financial interests.

## DATA AVAILABILITY

The cryo-EM maps of the mitoribosome have been deposited at the Electron Microscopy Data Bank (EMDB) with accession codes XXXX. Corresponding atomic models have been deposited in the Protein Data Bank (PDB) with PDB codes XXXX. The mass spectrometry proteomics data have been deposited to the ProteomeXchange Consortium via the PRIDE partner repository with the dataset identifier PXD018981.

## ONLINE METHODS

### Purification of *T. cruzi* and *L. tarentolae* mitochondria

For mitochondria purification, protocols adapted from Hauser et al. 1996^42^ and Schneider et al. 2007^43^ were used. Quickly, cells were re-suspended in SoTE buffer (600 mM sorbitol, 20 mM Tris-HCl pH 7.5, 1 mM EDTA) to a final concentration of 2.5×10^9^ cells/ml and lysed by nitrogen cavitation at 80 bar for 1 hour. After pressure release the lysate was centrifuged and resuspended in SoTM buffer (600 mM sorbitol, 20 mM Tris-HCl pH 7.5, 5 mM MgCl_2_) for DNase I treatment during 15 minutes. EDTA was added to stop DNAse treatment and the sample was centrifuged, resuspended in 2 volumes of SoTE containing 50% Histodenz™ (Sigma Aldrich D2158). For *L. tarentolae* the fraction was loaded onto a 21.7%/25%/28.3/31.6% Histodenz™ step gradients. Gradients were centrifuged for 45 minutes at 100,000 g and the mitochondrial fraction was collected at the 25%/28.3% interface. For *T. brucei* the fraction was loaded onto a 18%/21%/25%/28% Histodenz™ step gradient and the mitochondrial fraction was collected at the 21%/25% interface.

### Purification of mitochondrial ribosomes

For mitoribosome purification, protocols adapted from Waltz & Soufari et al.^10^ were applied. Quickly, mitochondria were re-suspended in Lysis buffer (20 mM HEPES-KOH, pH 7.6, 100 mM KCl, 20 mM MgCl_2_, 1 mM DTT, 1 % Triton X-100, 2 % DDM, supplemented with proteases inhibitors (Complete EDTA-free) to a concentration of 1 mg/mL and incubated for 15 min in 4°C. Lysate was clarified by centrifugation at 30.000 g, 20 min at 4°C. The supernatant was loaded on a 40% sucrose cushion in Monosome buffer (same as lysis buffer without Triton X-100 and with 0.02% DDM) and centrifuged at 235.000 g, 3h, 4°C. The crude ribosomes pellet was re-suspended in Monosome buffer and loaded on a 10-30% sucrose gradient in the same buffer and run for 16 h at 65,000 g. Fractions corresponding to mitoribosomes were collected, pelleted and re-suspended in Monosome buffer.

### Grid preparation

4 μL of the samples at a concentration of 2 μg/μl was applied onto Quantifoil R2/2 300-mesh holey carbon grid, which had been coated with thin home-made continuous carbon film and glow-discharged. The sample was incubated on the grid for 30 sec and then blotted with filter paper for 2 sec in a temperature and humidity controlled Vitrobot Mark IV (T = 4°C, humidity 100%, blot force 5) followed by vitrification in liquid ethane pre-cooled by liquid nitrogen.

### Single particle cryo-electron microscopy data collection

For the two data-sets (Full and dissociated complexes), data collection was performed on a Talos Arctica instrument (Thermofisher) at 200 kV using the EPU software (Thermofisher) for automated data acquisition. Data were collected at a nominal underfocus of −0.5 to −2.7 μm at a magnification of 120,000 X yielding a pixel size of 1.24 Å. Micrographs were recorded as movie stack on a Falcon III direct electron detector (Thermofisher), each movie stack were fractionated into 20 frames for a total exposure of 1 sec corresponding to an electron dose of 60 ē/Å2.

### Electron microscopy image processing

Drift and gain correction and dose weighting were performed using MotionCor2^44^. A dose weighted average image of the whole stack was used to determine the contrast transfer function with the software Gctf^45^. The following process has been achieved using RELION 3.0^46^. Particles were picked using a Laplacian of gaussian function (min diameter 280 Å, max diameter 480 Å). *L. tarentolae* full mitoribosome, after 2D classification, 303,140 particles were extracted with a box size of 400 pixels and binned four fold for 3D classification into six classes. Three classes depicting high-resolution features, consisting of 82,060 particles, were selected for refinement, yielding 3.9 Å resolution. After focused refinement with a mask on the LSU, the body and the head of the SSU, 3.6, 4 and 3.8 Å resolution were respectively obtained. For the assembly intermediate of the LSU, after 2D classification, 215,120 particles were extracted with a box size of 400 pixels and binned three fold for 3D classification into 8 classes. Two classes, consisting of 59,200 particles, depicting high-resolution features were selected for refinement, yielding 3.4 Å resolution.

For the *T. cruzi* full mitoribosomes reconstruction, 126,340 particles were selected after 3D classification for refinement, resulting in a 6 Å resolution reconstruction, which was further refine to 3.7 Å and 4.5 Å resolution for the LSU and SSU respectively. For the *T. cruzi* mt-IF3 SSU complex, 148,180 particles were selected after 3D classification for refinement, resulting in a 3.5 Å resolution reconstruction, which was further refine to 3.1 Å and 3.2 Å resolution for the SSU body and SSU head respectively. Determination of the local resolution of the final density map was performed using ResMap^40^.

### Structure building and model refinement

The atomic models of the kinetoplastid mitoribosomes were built into the high-resolution maps using Coot, Phoenix and Chimera. Atomic models from *E. coli* ribosome (5kcr)^47^, and trypanosoma mitoribosome (6hiv)^3^ were used as starting points for protein identification and modelisation. For the Leishmania mitoribosome, atomic models were build using *L. major* protein sequences due to the poor quality of *L. tarentolae* available data. The online SWISS-MODEL service was used to generate initial models for bacterial and mitochondria conserved r-proteins. Models were then rigid body fitted to the density in Chimera^48^ and all subsequent modeling was done in Coot^49^. For the LSU ribosomal RNA, the 23S rRNA from *E.coli* was docked into the maps and used as template from positioning and reconstruction. Punctual differences were done in Chimera using the “swapna” command line and the model was then further refined in Coot. The global atomic model was refined with VMD using the Molecular Dynamic Flexible Fitting (MDFF) then with PHENIX using a combination of real and reciprocal space refinement for proteins.

### Proteomic and statistical analyses of mitochondrial ribosome composition

Mass spectrometry analyses of the ribosome fractions were performed at the Strasbourg-Esplanade proteomic platform and performed as previously^10,50^. In brief, proteins were trypsin digested and mass spectrometry analyses were carried out by nano LC-ESI-MS/MS analysis on a Thermo QExactive+ mass spectrometer. Quantitative label-free proteomics analysis was performed through in-house bioinformatics pipelines.

### Figure preparation

Figures featuring cryo-EM densities as well as atomic models were visualized with UCSF ChimeraX^51^ and Chimera^48^.

---

**Extended Data Figure 1.**
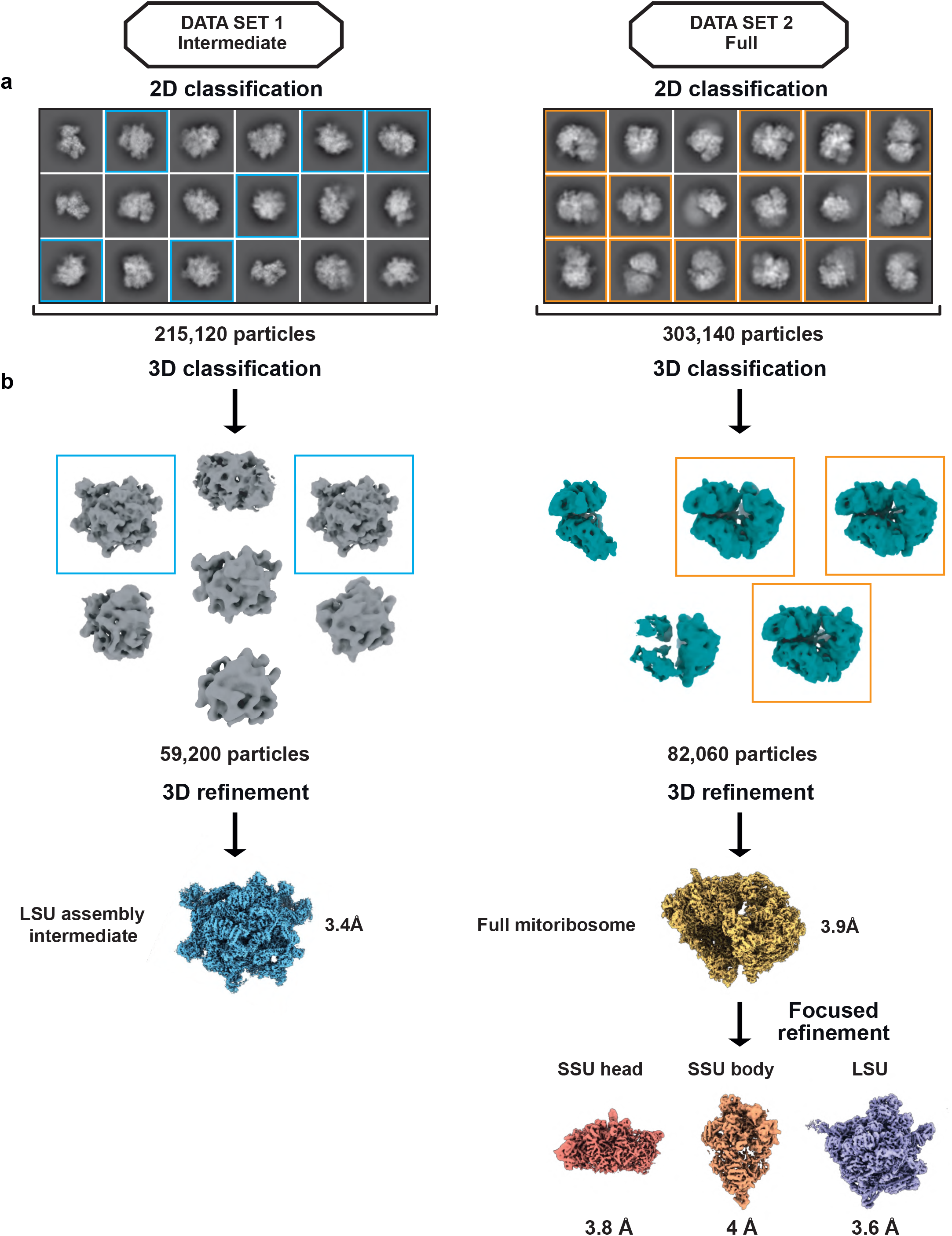
Data processing workflow of *L. tarentolae* data sets. Graphical summary of the processing workflow described in Methods for the *L. tarentolae* samples, resulting in the LSU assembly intermediate for Data set 1 and the full mitoribosome for Data set 2. 2D classes are presented in (**a**) and 3D processing and refinement presented in (**b**).

**Extended Data Figure 2.**
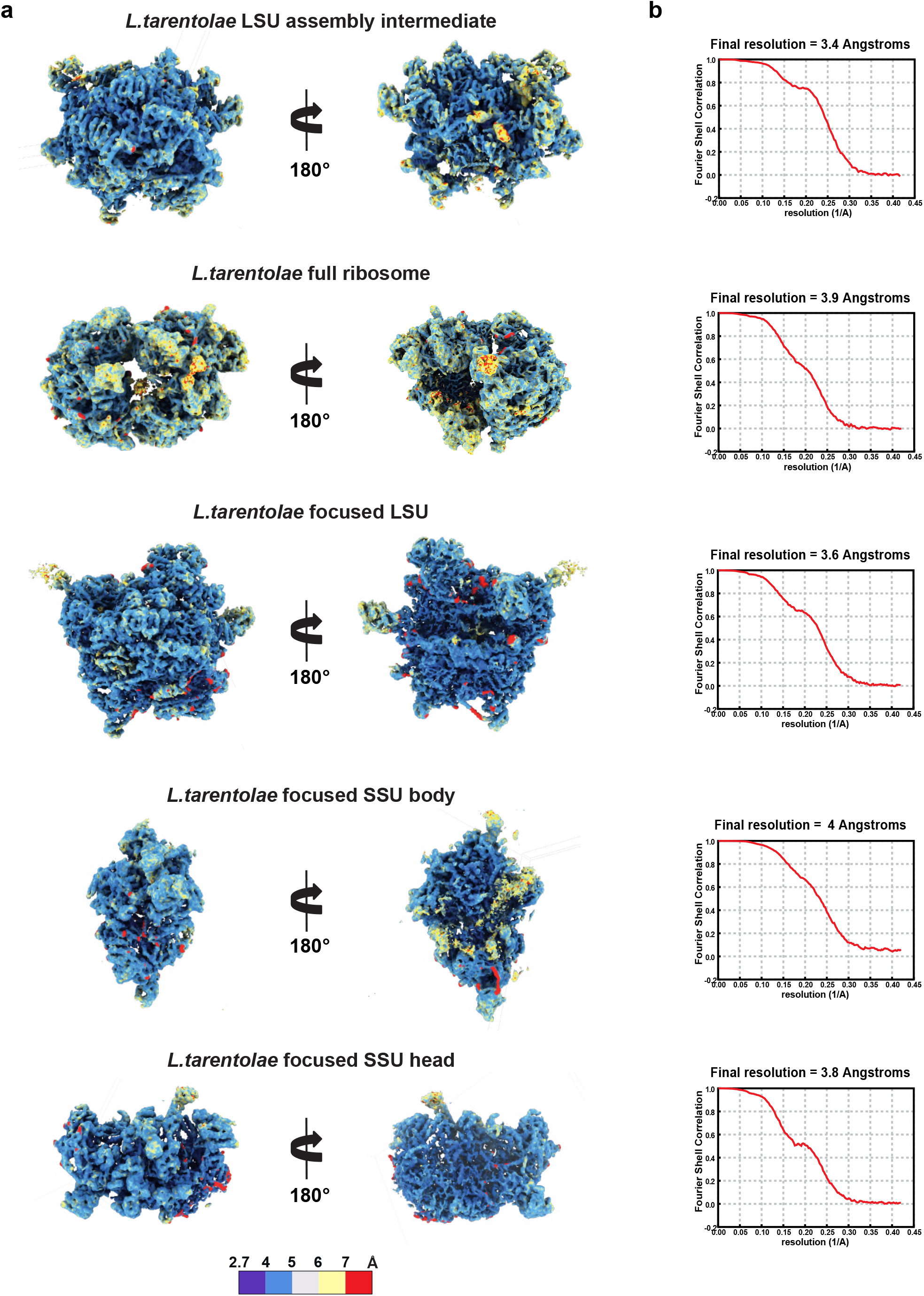
Local resolution of *L. tarentolae* final reconstructions. Local resolutions of the different reconstructions are presented. The maps are colored by resolution, generated using ResMap^40^ (**a**). For both reconstructions FSC plots are displayed for resolution estimation (**b**). The map resolution is calculated at the 0.143 threshold.

**Extended Data Figure 3.**
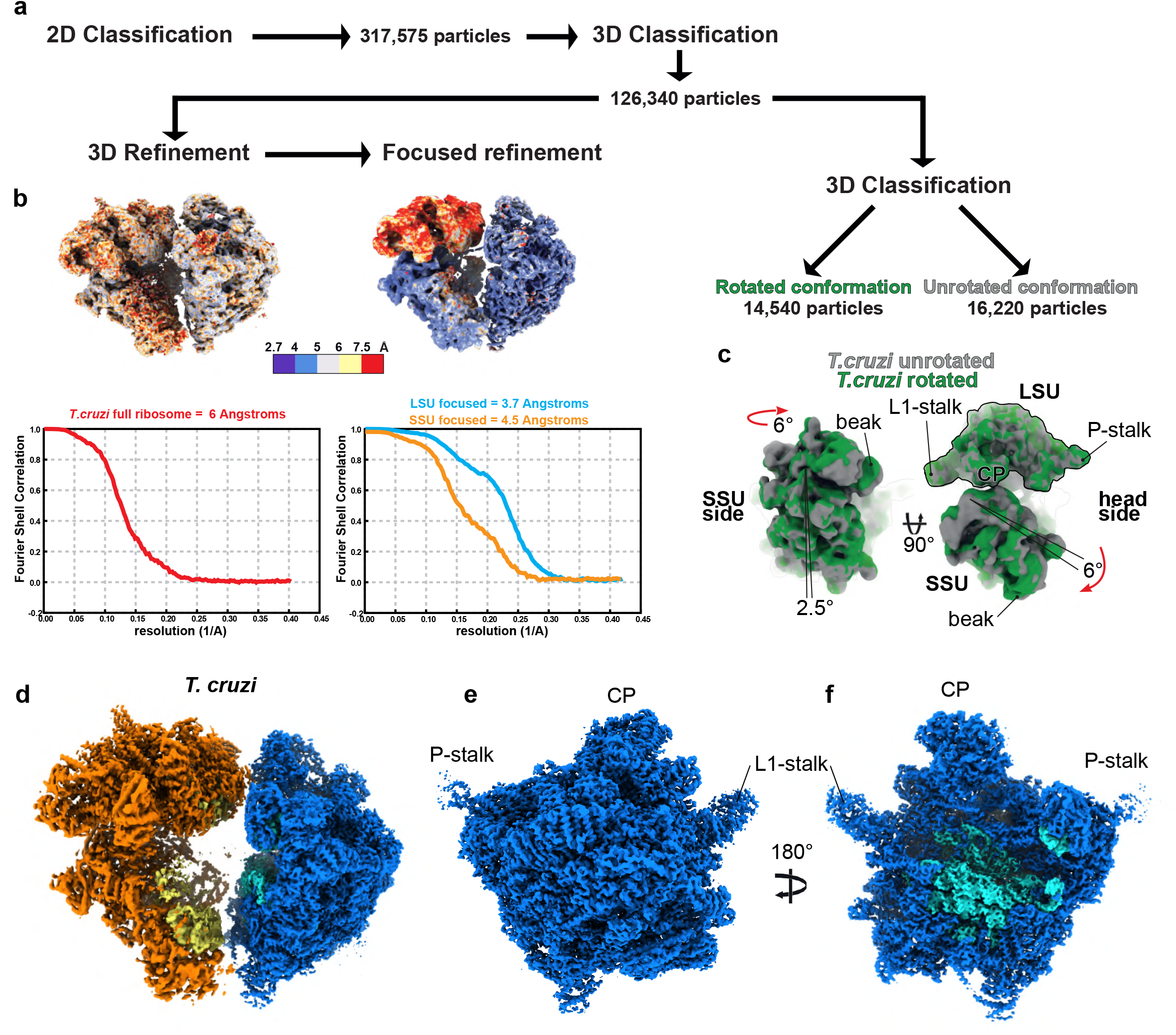
Data processing and cryo-EM reconstructions of *T. cruzi* mitoribosome. Graphical summary of the processing workflow *T. cruzi* full mitoribosome. 2D classification and 3D processing are presented in (**a**) with respective ResMap^40^ representation and FSC plots presented in (**b**). Second round of 3D classification lead to the identification of two rotational states presented in (**c**). Cryo-EM reconstructions of *T. cruzi* full mitoribosome (**d**) as well as the mature LSU, seen from the solvent view (**e**) and the interface view (**f**). Compared with the *L.tarentolae* reconstructions presented in Figure 1, the ribosomes are nearly identical.

**Extended Data Figure 4.**
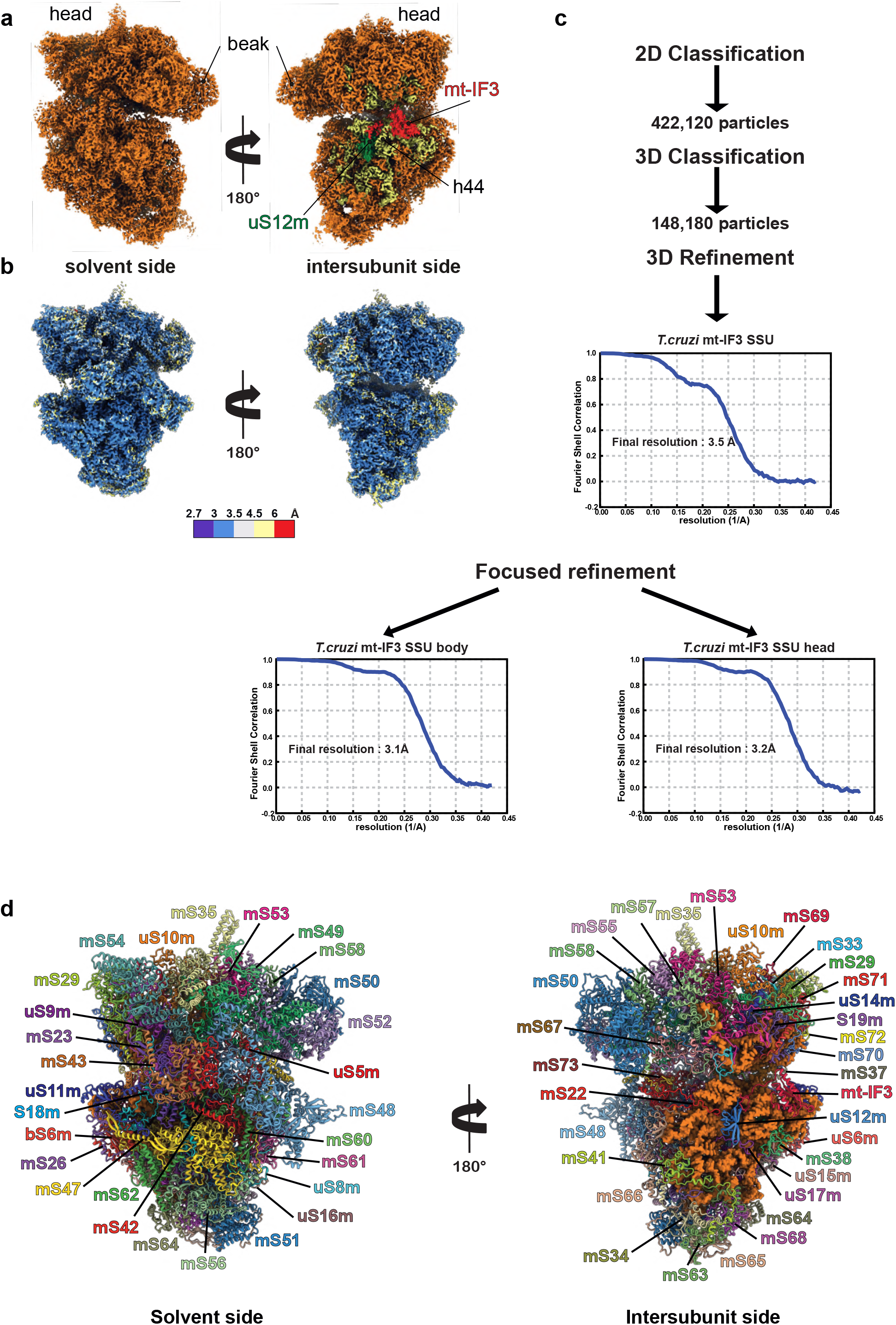
Data processing and model of the mt-IF3 SSU of *T. cruzi*. Cryo-EM map reconstruction of the *T. cruzi* in presence of mt-IF3 is shown in (**a**), with r-proteins colored in orange, 9S rRNA in yellow, mt-IF3 in red and uS12m in green. Local resolution generated using ResMap^40^ are shown in (**b**), and the processing workflow with the respective FSC plots for initial and focused 3D refinement are displayed in (**c**). The resulting atomic model is presented in (**d**), each r-proteins are individually annotated, 9S rRNA is colored in orange.

**Extended Data Figure 5.**
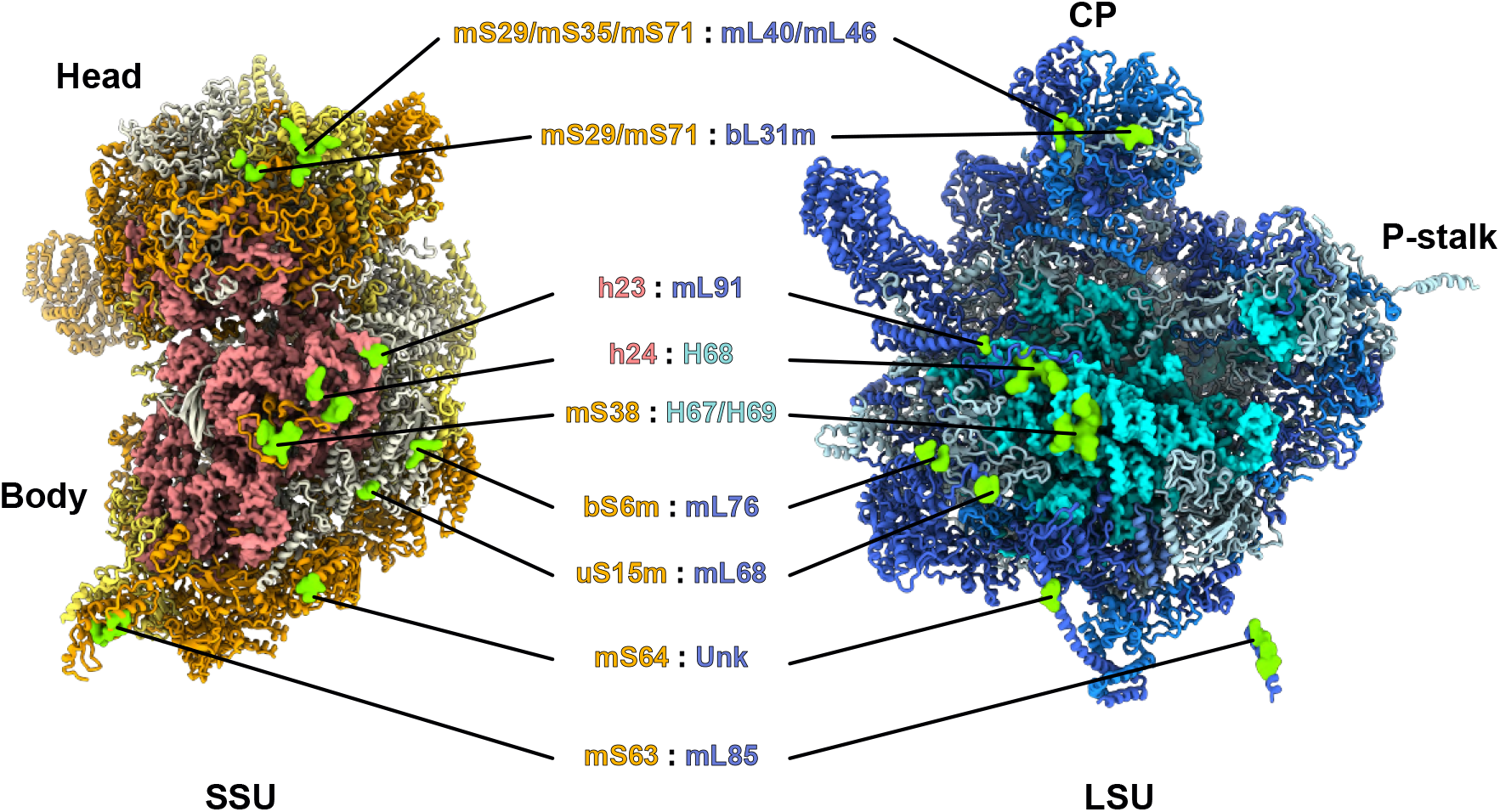
Intersubunit bridges. List of the intersubunit bridges observed in the mature and full kinetoplastid mitoribosomes. Both the SSU and LSU are presented on the intersubunit side and bridge positions are indicated in green. Most of the bridges involve protein:protein interactions, except for the highly conserved rRNA:rRNA bridge between h24 and H68. Overall, the other bridge positions seem to be common with other mitoribosomes and bacterial ribosomes but involve mitochondria and kinetoplastid-specific r-proteins, for example, bL31m is the only bacterial-type r-protein involved in the bridges between the SSU’s head and LSU’s CP. One truly kinetoplastid-specific bridge is observed with mS63 and mL85, which involves kinetoplastid only r-proteins.

**Extended Data Figure 6.**
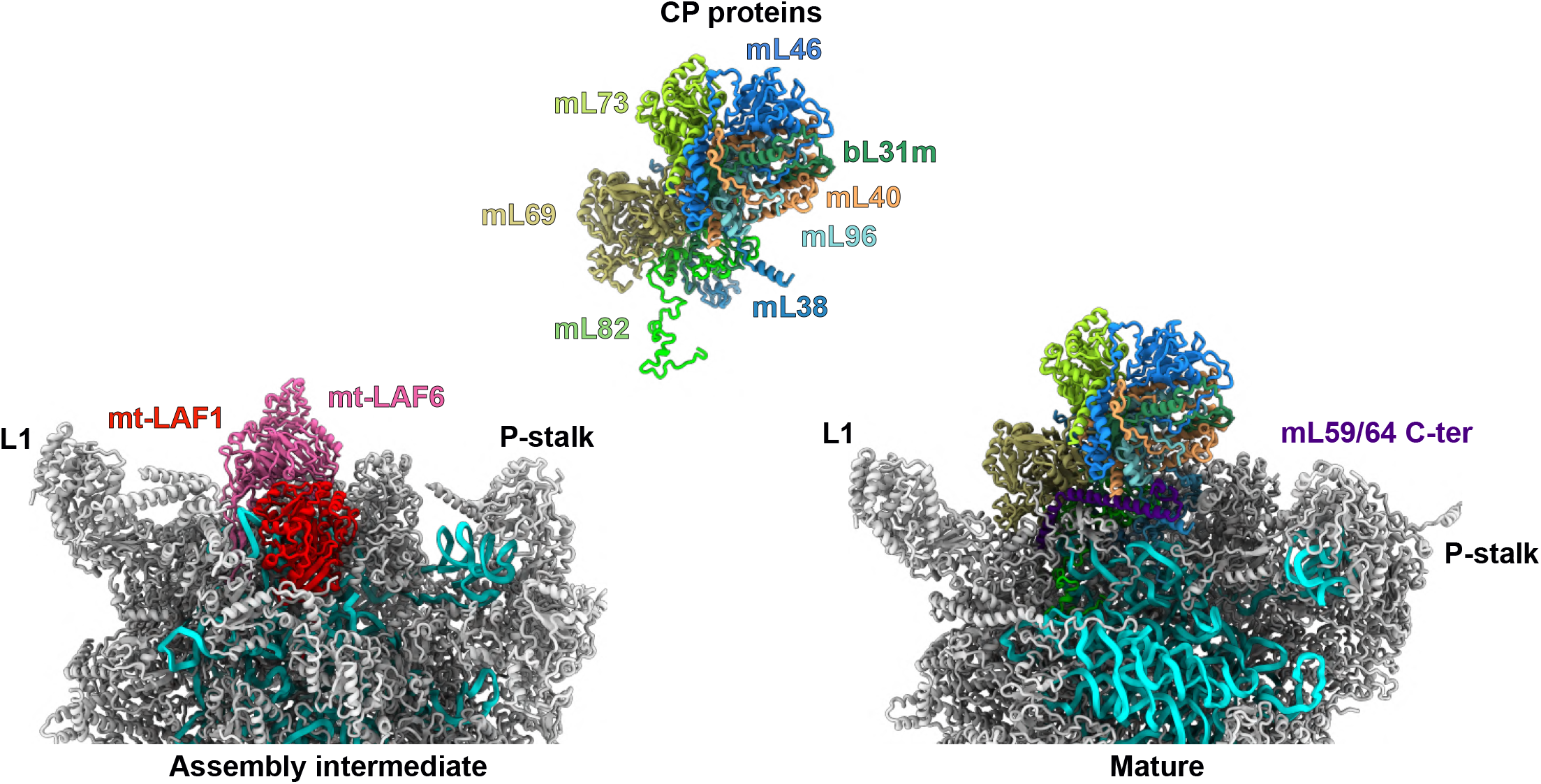
Central protuberance maturation in Leishmania LSU. Comparison of the assembly intermediate and mature CP area of the LSU. In the assembly intermediate, CP proteins are prevented from binding by mt-LAF1 and mt-LAF6. Upon release of the maturation factors the whole CP block can dock the LSU and additional r-proteins contacting the CP, like bL33m and the C-terminus of mL59/64 can lock the structure. 12S rRNA is displayed in cyan.

**Extended Data Figure 7.**
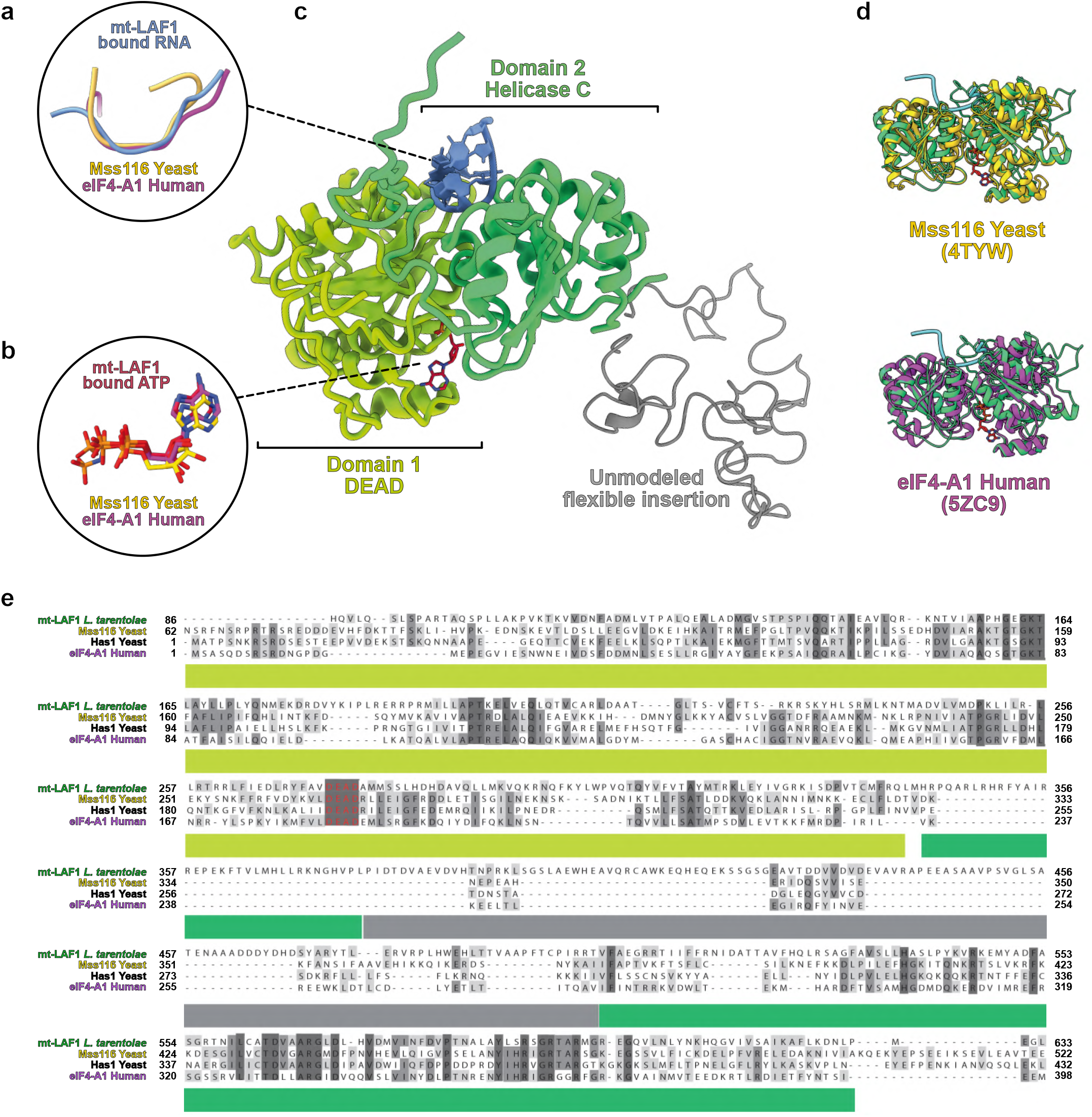
Structural conservation and sequence comparison of the DEAD-box RNA helicase mt-LAF1. mt-LAF1 is shown in (**c**) with its DEAD and helicase domains colored in green and its unmodeled flexible insertion shown in grey. Aside the insertion, the helicase appears to be conserved with enzymes of the DEAD-box family, highlighted by the superimpositions (**d**) and sequence alignments (**e**) of mt-LAF1 with yeast mitochondrial Mss116 enzyme^12,13^, or cytosolic Has1 and eIF4-A1. The enzyme is in closed state (**d**) and the interaction with RNA (**a**) as well as with the ATP ligand (**b**) is highly conserved.

**Extended Data Figure 8.**
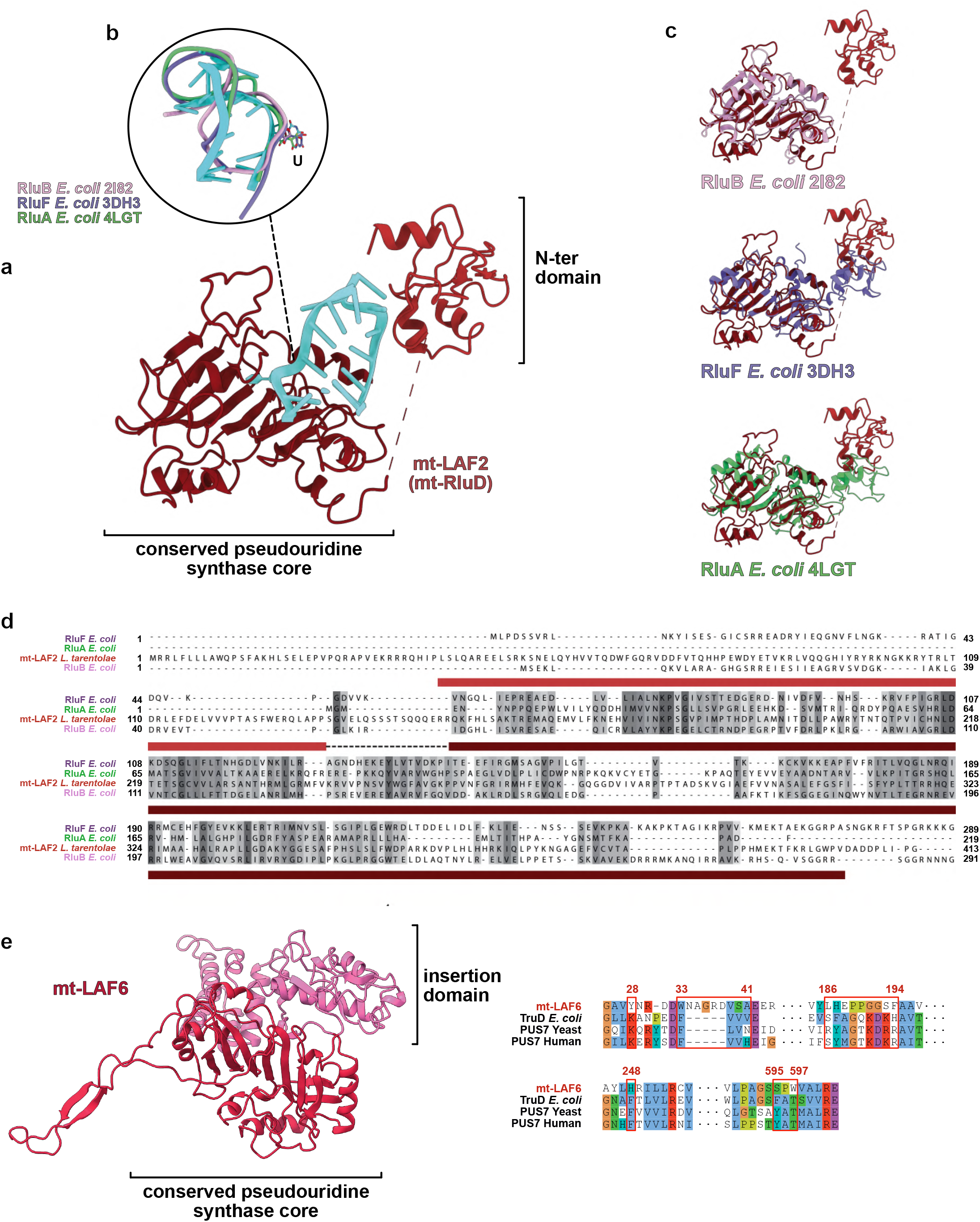
Analysis of the pseudouridine synthases mt-LAF2 and mt-LAF6. Structural conservation and sequence comparison of the two pseudouridine synthases mt-LAF2 and mt-LAF6 are shown. In (**a**) the pseudouridine synthase mt-LAF6, homolog of the bacterial enzyme RluD^19,20^, is shown with its rRNA target (blue). The enzyme is particularly conserved with enzymes of the Rlu family, as highlighted by the superimpositions (**c**) and sequence alignments (**d**) of mt-LAF2 with *E. coli* Rlu enzymes. Its mode of action is also conserved, as shown in (**b**). In (**e**) the analysis of mt-LAF6 shows the presence of an insertion domain (pink) and sequence alignment with other pseudouridine synthases highlight that none of the catalytic sites of mt-LAF6 are conserved, mt-LAF6 is thus inactive.

**Extended Data Figure 9.**
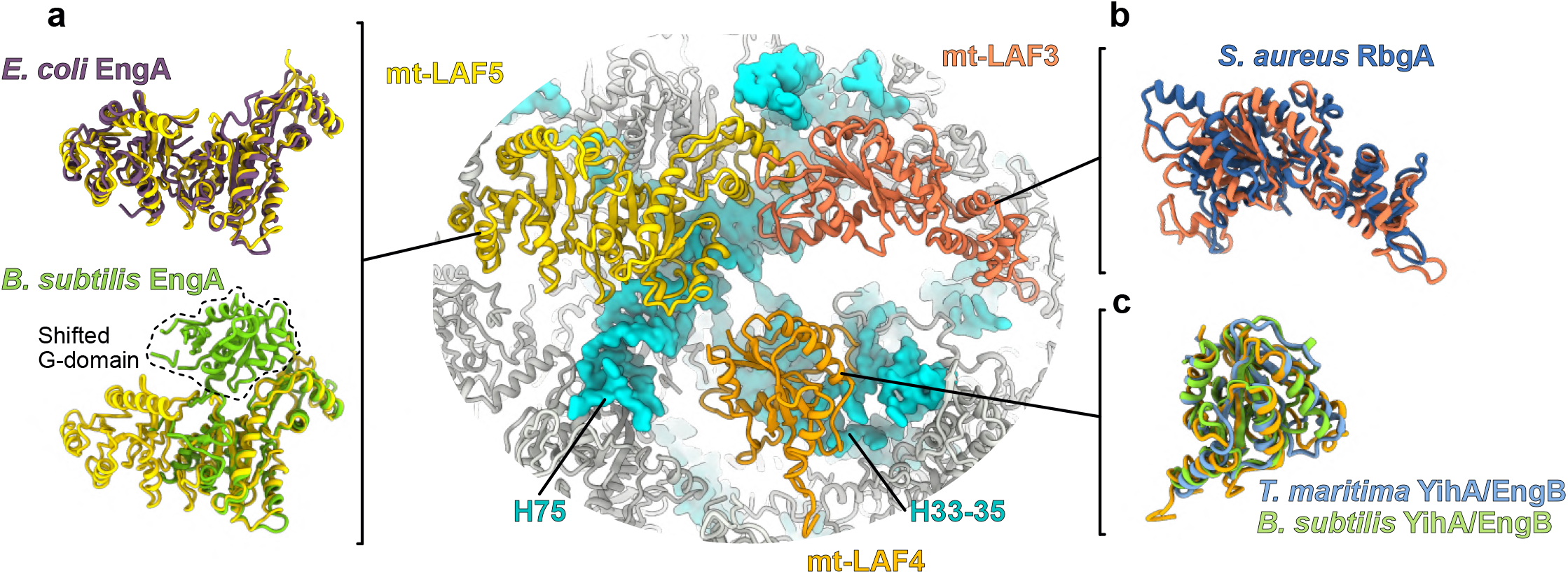
Conservation of the three GTPases with their bacterial homologs. The three GTPases mt-LAF3, mt-LAF4 and mt-LAF5 are compared with their bacterial homologs. In (**a**) mt-LAF5 is compared with *B. subtilis* EngA crystal structure obtained with GDP, and *E. coli* EngA structure obtained by cryo-EM in complex with the LSU with excess of GMP-PNP, a non-hydrolysable analog of GTP (PDB: 3J8G and 5M7H). The *B. subtilis* one has its first G-domain shifted compared to *E. coli* EngA and mt-LAF5 which represent one of the alternative structural state of the GTPase. Thus, mt-LAF5 is in GTP state. (**b**) shows mt-LAF3 superimposed with *S. aureus* RbgA(PDB: 6G14). In (**c**) mt-LAF4 is superimposed with two crystal structures of its bacterial homolog YihA/ EngB (PDB: 3PQC and 1SVW). This protein was never observed in native assembly, but our reconstruction appear to be quasi-identical to the crystal structures previously obtained. In context of the LSU assembly intermediate, it interacts with H54 and appears here to block uL2m association.

**Extended Data Figure 10.**
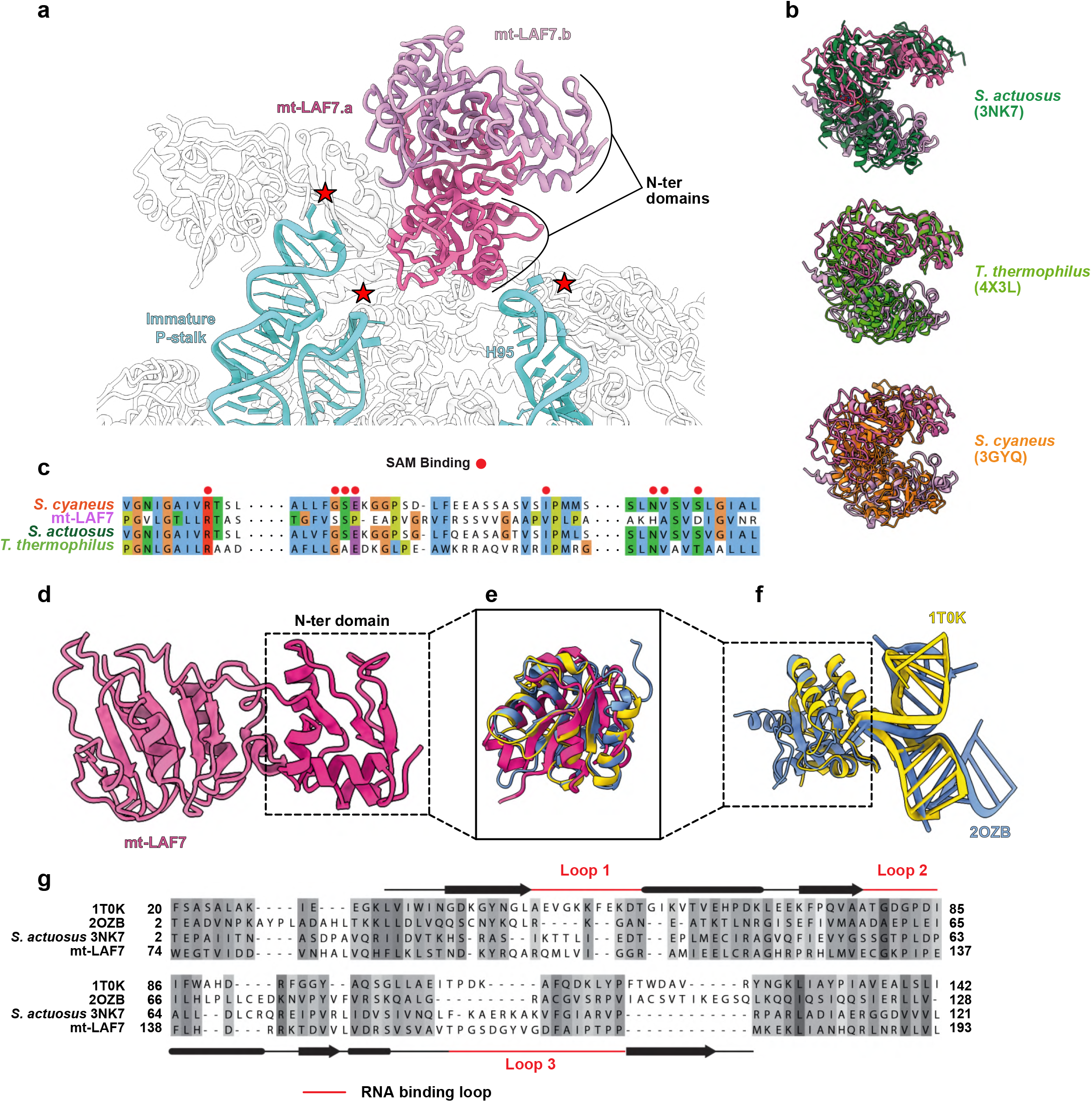
Analysis of the RNA2’O-methyltransferase mt-LAF7. The homodimeric mt-LAF7.a/mt-LAF7.b 2’O-methyltransferases are shown in the assembly intermediate context (**a**), with nearby possible rRNA targets - immature P-stalk and H95 - highlighted with a red star. Structural conservation is highlighted by structural superimpositions with several bacterial homologs, notably with *S. actuosus* NHR involved in P-stalk methylation^41^ (**b**), and crucial residues for the activity involved in S-adenosyl-l-methionine (SAM) binding are shown in (**c**). In (**d-g**) an analysis of the RNA binding N-terminal domain is shown. The N-terminal domain of mt-LAF7 (**d**) is superimposed (**e**) and the sequences are compared (**g**) with two eukaryotic proteins harbouring a similar dsRNA binding domain, which were structurally characterized in complex with their double stranded RNA target (**f**) as well as with *S. actuosus* NHR domain.

## Notes

### Competing Interest Statement

The authors have declared no competing interest.

### Summary of Updates

Text was extended from letter format to full article, and several figures were added.

